# Post-translational modification of SPATULA by SECRET AGENT and SPINDLY promotes organ symmetry transition at the gynoecium apex

**DOI:** 10.1101/2023.04.28.538690

**Authors:** Yuxiang Jiang, Seamus Curran-French, Samuel W.H. Koh, Iqra Jamil, Luca Argirò, Sergio Lopez, Carlo Martins, Gerhard Saalbach, Laila Moubayidin

## Abstract

The establishment of organ symmetry during multicellular development is a fundamental process shared by most living organisms. Here, we investigated how two *O*-glycosyltransferases of *Arabidopsis thaliana*, SPINDLY (SPY) and SECRET AGENT (SEC) synergistically promote a rare bilateral-to-radial symmetry transition during patterning of the plant reproductive organ, the gynoecium. SPY and SEC modify N-terminal residues of the bHLH transcription factor SPATULA (SPT) *in vivo* and *in vitro* by attaching *O*-fucose and *O*-linked-β-N-Acetylglucosamine (*O*-GlcNAc), to promote style development. This post-translational regulation does not impact SPT homo- and hetero-dimerisation events with INDEHISCENT (IND) and HECATE 1 (HEC1), although it enhances the affinity of SPT for the kinase *PINOID* (*PID*) gene locus to promote transcriptional repression. Our findings reveal a previously unrecognized mechanism for *O*-GlcNAc and *O*-fucose post-translational decorations in controlling style development and offer the first molecular example of a synergistic role for SEC and SPY in plant post-embryonic organ patterning.

## MAIN

A major challenge during morphogenesis of plant and animal organs is establishing a ground symmetry type, *i.e.,* implementing radial or bilateral symmetry in concert with polarity axes and tissue specification. The symmetry type that an organ/organism adopts during development is inextricably connected to its function and ultimately contributes to the fitness and evolution of a species^1–3^. For example, in plants the geometric arrangement of flower organs is coordinated with the specification of the body axes to guide the morphogenesis of radial (actinomorphy) and bilateral symmetric flowers^4,5^. Within the flower itself a sophisticated organ known as the gynoecium ensures fertilization and seed production. Depending on plant species and the developmental window gynoecium shape and symmetry type may vary to complement its function^6,7^.

Despite the importance of the gynoecium for fitness and seed production, the underlying molecular mechanisms underpinning dynamic symmetry establishment during development remain unelucidated.

The gynoecium of *Arabidopsis thaliana* undergoes a rare bilateral-to-radial symmetry transition during organogenesis^8,9^. During development, the apical end of the bilaterally symmetric ovary adopts radial symmetry forming a compact cylindrical structure known as the style^8^. This symmetry transition requires dynamic control of auxin distribution (via biosynthesis^10,11^, signalling^12,13^ and transport^8^) as well as crosstalk with the antagonistic proliferating signal mediated by the hormone cytokinin (CK)^14,15^ This is orchestrated by a set of transcription factors (TFs)^16^ interacting at the transcriptional and protein level^17^.

Among these TFs, the activity of SPATULA (SPT) - a key regulator of medial tissue identity and symmetry transition during gynoecium development^18^ – is pivotal in orchestrating auxin accumulation at the apical-medial cells^8^ as well as coordinating the medio-lateral and adaxial-abaxial polarity axis^8,9,19^ and repressing CK-mediated cell-proliferation input at the gynoecium apex^20^. SPT forms homo- and hetero-dimers with specific TF partners to modulate several aspects of style development^16,17,21–23^. Accordingly, *SPT* loss-of-function mutants fail to develop a radial style at the gynoecium apex^18^ (producing a so-called split-style) and this phenotype is exacerbated by mutations in other basic Helix-loop-Helix (bHLH) TFs such as IND^24^ and HECs^9,16^.as well as members of the NGATHA protein family^17^.

Because of its stable spatio-temporal expression within the apical-medial tissues during gynoecium development^18^ we hypothesised that the dynamic activity of SPT required for style patterning may be regulated at the protein level. Interestingly, a proteomic study showed that SPT is post-translationally modified by *O*-linked-β-N-Acetylglucosamine (*O*-GlcNAc)^25^. *O*-GlcNAcylation has been associated with stem-cell survival, embryo development and *HOX* gene expression during body-axis formation in animals^26–28^.

In Arabidopsis, the enzyme SECRET AGENT (SEC) catalyses the addition of *O*-GlcNAc from UDP-GlcNAc to serine (Ser) and/or threonine (Thr) residues of acceptor-substrate proteins^29^. A functionally related enzyme encoded by the gene locus *SPINDLY* (*SPY*)^30^ functions as an *O*-fucosyltransferase (POFUT) attaching monofucose (*O*-fucose) to target substrates^30,31^. Notably, while single *sec* and *spy* mutant alleles are perfectly viable in *Arabidopsis*, *sec spy* double mutants are embryo lethal, highlighting the synergistic and fundamental importance of both enzymes for plant development^32^.

By means of genetic, molecular, biochemical and proteomic experiments, we herein demonstrate a role for SEC and SPY in style development and radial symmetry establishment via post-translational regulation of SPT activity. We demonstrate that SPT directly interacts with SEC and SPY, which modify the N-terminus of SPT by adding *O*-GlcNAc and *O*-fucose respectively, both *in vivo* and *in vitro*. Moreover, via a genetic complementation assay we provide evidence that specific modified residues located in two N-terminal peptides, are essential for SPT function *in vivo,* accounting for radial symmetry establishment. Furthermore, we show both enzymes enhance the transcriptional activity (as well as affinity) of SPT for *PINOID* (*PID*)^33^ but do not impact SPT nuclear localization, protein stability or dimerization events with itself, IND or HEC1. Accordingly, genetic epistasis analysis and exogenous CK treatments, corroborate a model in which SPT activity is primed by SEC and SPY enzymatic activity to promote style development by fine-tuning the auxin/cytokinin balance.

Our data characterise the first post-embryonic target modified synergistically by both SEC and SPY during plant organ development and highlights how the function of these sugar-based PTMs are at the crux of tissue-identity, body-axes formation, and the genetic network downstream of key organ regulators important for organ symmetry in plants.

## RESULTS

### SPT is post-translationally modified by *O*-GlcNAc and *O*-fucose *in vivo* and *in vitro* via the activity of SEC and SPY transferases

To understand whether a post-translation mechanism based on *O*-glycosylation could regulate the SPT-mediated control of radial organ symmetry establishment, we performed Higher energy Collisional Dissociation (HCD) fragmentation Mass Spectrometry (MS/MS) analysis on nuclear extracts from *pSPT:SPT:YFP/spt-12* complementation line inflorescences (Extended Data Fig. 1a,b). This analysis identified specific Ser and Thr residues at the N-terminus of SPT were modified by two *O*-linked sugar moieties, *O*-fucose and *O*-GlcNAc. *In vivo*, Ser23, Ser24 (recovered in peptide_1) and Ser60, Ser61 (recovered in peptide_2) hosted both sugar groups, while Thr43 (peptide_1) and Ser65, Ser68, Thr71, Thr73 (peptide_2) were modified specifically by *O*-GlcNAc (Fig. 1a,b, Extended Data Fig. 1c). Whilst Ser25 (peptide_1) was modified solely by *O*-fucose (Fig. 1a, Extended Data Fig. 1c). The percentage of both modifications on the two N-terminal peptides of SPT were then quantified. Levels of *O*-fucose modification were higher compared with *O*-GlcNAc on peptide_1, while on peptide_2 *O*-GlcNAcylation was conspicuously elevated and *O*-fucosylation was barely detected (Fig. 1d).

**Fig. 1.**
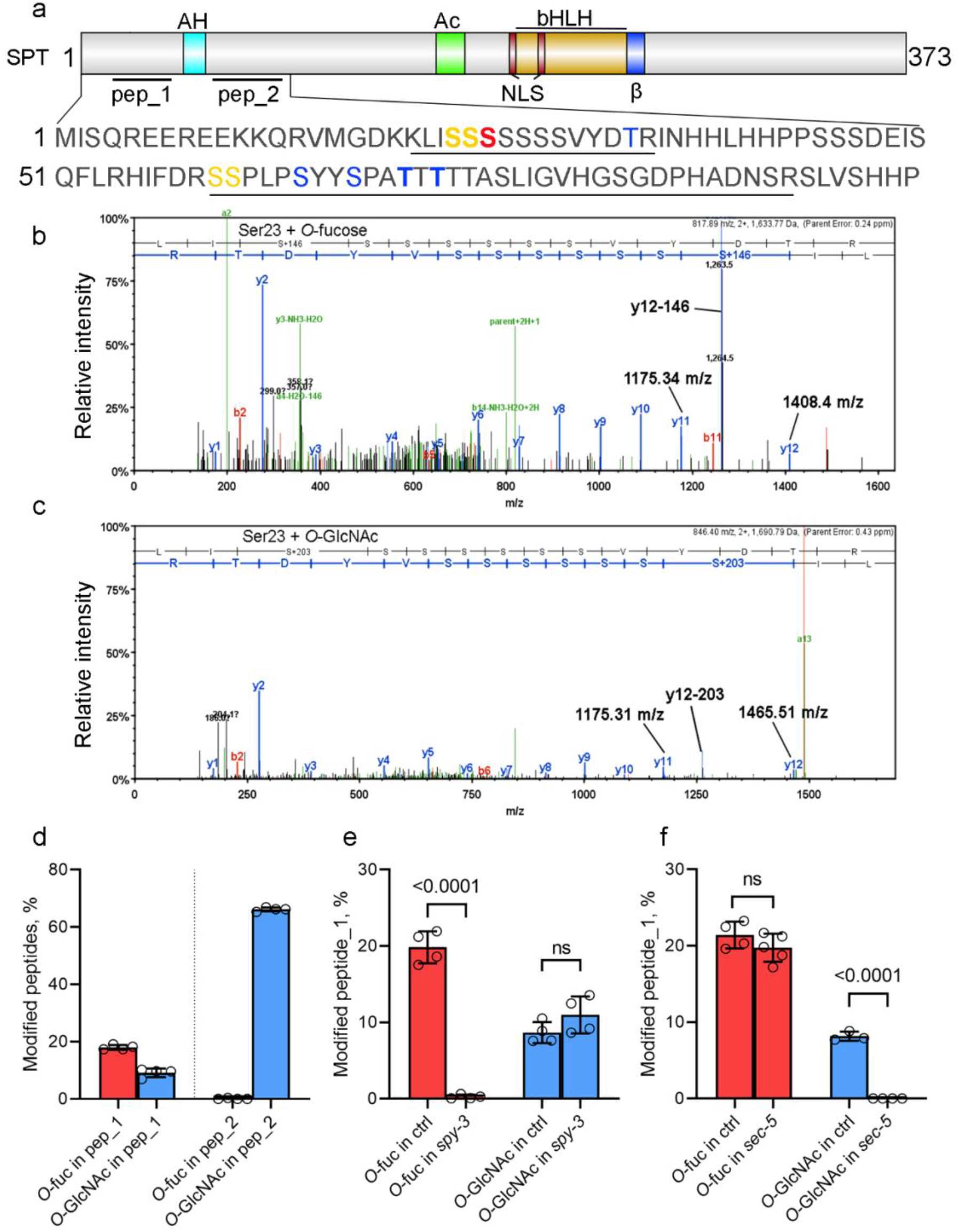
SPT is modified *in vivo* by both *O-*fucose and *O-*GlcNAc via SPY and SEC activity respectively. **a**, Illustration of SPT full-length protein and its domains (AH, Amphipathic helix; Ac, Acidic helix; NLS, nuclear localization signal; bHLH basic Helix-loop-Helix domain, ß beta-strand^23^) The position of the two peptides (pep_1 and pep_2) targeted by *O-*glycosyl PTM is displayed flanking the AH domain. The SPT N-terminal amino acid sequence is displayed, including pep_1 and pep_2 (underlined sequences). Modified residues are colour-coded as follows: Blue, *O-*GlcNAc; Red, *O-*fucose; Yellow, both modifications. S/T residues in bold were identified as frequently modified in the MS analysis. **b** and **c**, Spectra of SPT residue Ser23 modified by *O-*fucose (b) and *O-*GlcNAc (c). **d-f,** Quantification of the percentage of *O-* fucosylation (green bars) and *O-*GlcNacylation (orange bars) recovered on both SPT peptides in the *pSPT:SPT-YFP/spt* complementation line inflorescences (d), and quantification of the same modifications recovered on pep_1 in *spy-3* (e) and *sec-5* (f) mutant backgrounds. A minimum of three independent experiments were repeated for each genotype. Values shown are means±SD. Significant differences and P values are indicated in the graph following student’s *t* test.

To assess whether modifications of SPT by *O*-GlcNAc and *O*-fucose on SPT were dependent on SEC and SPY activity respectively, we crossed a *spt-12/pSPT:SPT:YFP* complementation line with a single loss-of-function mutant of *SEC* (*sec-5^34^*) and a catalytically redundant mutant of SPY (*spy-3^35^*) (Extended Data Fig. 2a). Using *spt-12sec-5/pSPT:SPT:YFP and spt-12spy-3/pSPT:SPT:YFP* inflorescences we performed a similar HCD MS/MS analysis and compared the percentage of modifications in relation to the *spt-12/pSPT:SPT:YFP* segregating controls. Since *O*-fucosylation was hardly detected on peptide_2 we focused this analysis on peptide_1. Our data showed that *O*-fucose modification was completely abolished on peptide_1 in the *spy-3* background, while *O*-GlcNAc was present at comparable levels (Fig. 1e). On the other hand, in the *spt-12sec-5/pSPT:SPT:YFP* line we did not observe any change in the percentage of detectable *O*-fucose while *O*-GlcNAc was no longer detected in this background (Fig.1f). This analysis demonstrates that SPY and SEC modify SPT by *O*-fucose and *O*-GlcNAc, respectively, targeting both shared and specific residues. Our analysis highlights that the enzymes do not compensate for the activity of each other in their respective mutant backgrounds since we did not observe changes in transcript levels of *SPY* and *SEC* in *sec-5* and *spy-3* inflorescences, respectively (Extended Data Fig. 2b), nor an increase in their enzymatic activity as assessed by PTMs recovered on SPT (Fig.1e,f).

To confirm that SEC and SPY directly modify SPT post-translationally by *O*-GlcNAc and *O*-fucose we performed *in vitro* enzymatic assays to test the ability of recombinant SEC (*5TPR-SEC*)^30^ and SPY (*3TPR-SPY*)^30^ to directly modify full-length SPT protein (6xHis-SPT) (Extended Data Fig. 3a) in the presence and absence of their specific donor substrate (UDP-GlcNAc and GDP-fucose). Collectively, we found that eight residues of SPT were modified by both *O*-GlcNAc and *O*-fucose, seven were modified specifically by *O*-fucose and two were modified solely by *O*-GlcNAc (Extended Data Fig. 3b) including modification of all Ser and Thr residues previously identified *in vivo* (Fig. 1a).

Furthermore, no differences were observed for *SPT* transcript levels from *spy-3* and *sec-5* inflorescences compare to the wild-type (Extended Data Fig. 2c). Moreover, no reduction in SPT stability (as measured by the intensity of the SPT-YFP signal) or alterations in its subcellular localization were observed in the single *spy-3* and *sec-5* mutants (Extended Data Fig. 2d,e). Altogether, the results corroborate the idea that SEC and SPY work upstream of SPT acting at the post-translational level.

Thus, our data proves that specific residues of SPT can host both *O*-GlcNAc and *O*-fucosyl moieties, which are catalyzed by SEC and SPY *in vivo* and *in vitro*.

### SPT interacts *in vivo* and *in vitro* with SEC and SPY

To support a role for SEC and SPY in controlling SPT at the post-translational level, we tested whether these enzymes could directly interact with SPT. To this end, we employed Co-Immunoprecipitation (Co-IP) and split luciferase assays in *N. benthamiana* (Fig. 2a,b) alongside yeast-two-hybrid (Y2H) experiments (Fig. 2c and Extended data Fig.4) which all demonstrated direct interactions between SPT-SPY and SPT-SEC full-length proteins. Moreover, Y2H experiments also showed that both SPY and SEC bind to SPT via their N-terminal tetratricopeptide repeats (TPRs)^30^, TPRs 1-11 of SPY and TPRs 1-13 of SEC, while the C-terminal catalytic domain showed no growth of the yeast cells on selective media (Extended data Fig.4).

**Fig. 2.**
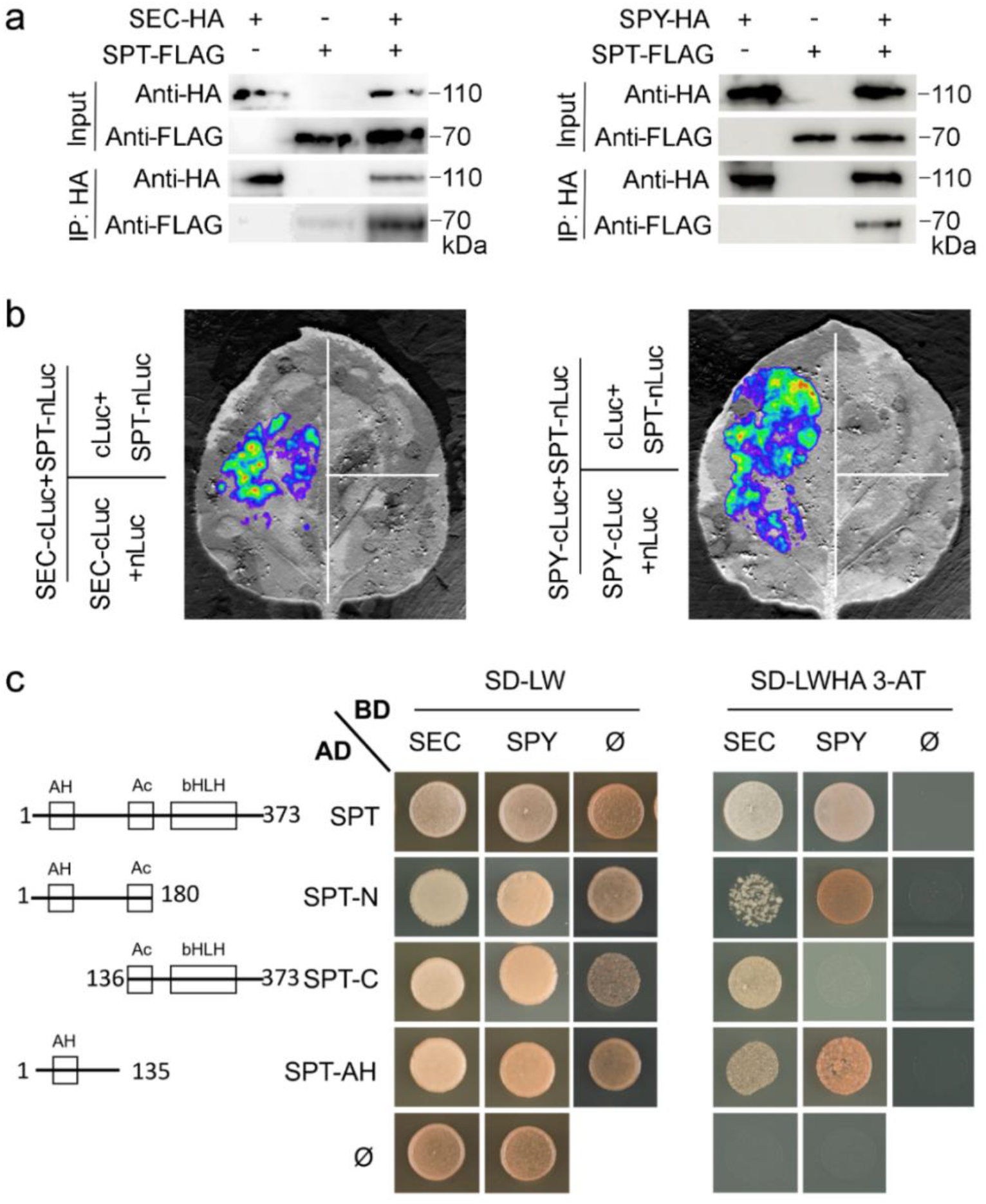
SPT directly interacts with SEC and SPY. **a**, Co-IP experiments performed in tobacco leaves showing the full-length SPT-FLAG protein co-immunoprecipitated with full-length SEC-HA (left panel) or SPY-HA (right panel) recombinant proteins. **b**, Split luciferase assay performed in tobacco leaves showing the interaction between full-length SPT-nLuc with both SEC-cLuc (left panel) and SPY-cLuc (right panel). **c**, Yeast-two-hybrid assay displaying SPT domains (left panel) sufficient for interactions with SPY and SEC full-length proteins. Positive co-transformants were selected on SD-WL (middle panel) and grown on SD-WLHA, 2.5 mM 3-AT (right panel) to determine strong interactions.

Interestingly, most of the residues modified on SPT were positioned at peptides flanking an Amphipathic (AH) helix at the N-terminus of SPT (Fig. 1A), a domain that supports the transcriptional activation activity of SPT^36^. Thus, to investigate the domains responsible for mediating the enzyme interactions with SPT, we performed Y2H experiments using the full-length sequences of SPY and SEC in combination with truncated versions of SPT. Our results showed the SPT N-terminal domain (spanning residues 1-180) -containing the Amphipathic and Acid (Ac) domains^36^- as well as a fragment containing only the Amphipathic domain (residues 1-135) - were both sufficient to produce positive interactions with SEC and SPY full-length proteins (Fig. 2c and Extended data Fig.4) but did not interact with HEC1 - a known interactor of SPT (Extended data Fig.4). On the other hand, the SPT C-terminal truncation (residues 136-373) including both the Ac and bHLH domains, did not interact with SPY but could interact with SEC as well as HEC1 (Fig. 2c and Extended data Fig.4). These results are in line with a working model in which binding of SPT with SPY occurs through the N-terminal domain of SPT containing the modified residues flanking the AH domain, whilst SEC interacts with the full-length SPT protein. These results also shed light on the signalling modulation occurring at the SPT N-term, which is important for transcriptional regulation and carpel patterning^36^.

### SEC and SPY play positive roles in style development and radial symmetry establishment

If the activity of *Arabidopsis* SEC and SPY control style development at the gynoecium apex, defects in either style specification or a break in radial symmetry should be observed in mutants for *sec* and *spy*. Despite decades of genetic analysis on single mutants for these two enzymes such defects have ever been reported^32,37,38^. Our scanning electron microscopy (SEM) analysis of loss-of-function *spy-4^38^*and *sec-5* single mutant gynoecia, as well as the catalytically defective mutants *spy-3* and *sec-2^37^* did not show any significant defects in style development (Fig. 3a). The radial style observed in *sec* and *spy* mutants is also in line with normal *SPT* expression within the style of *sec*/*spy* mutants (Extended data Fig. 2c,d).

**Fig. 3.**
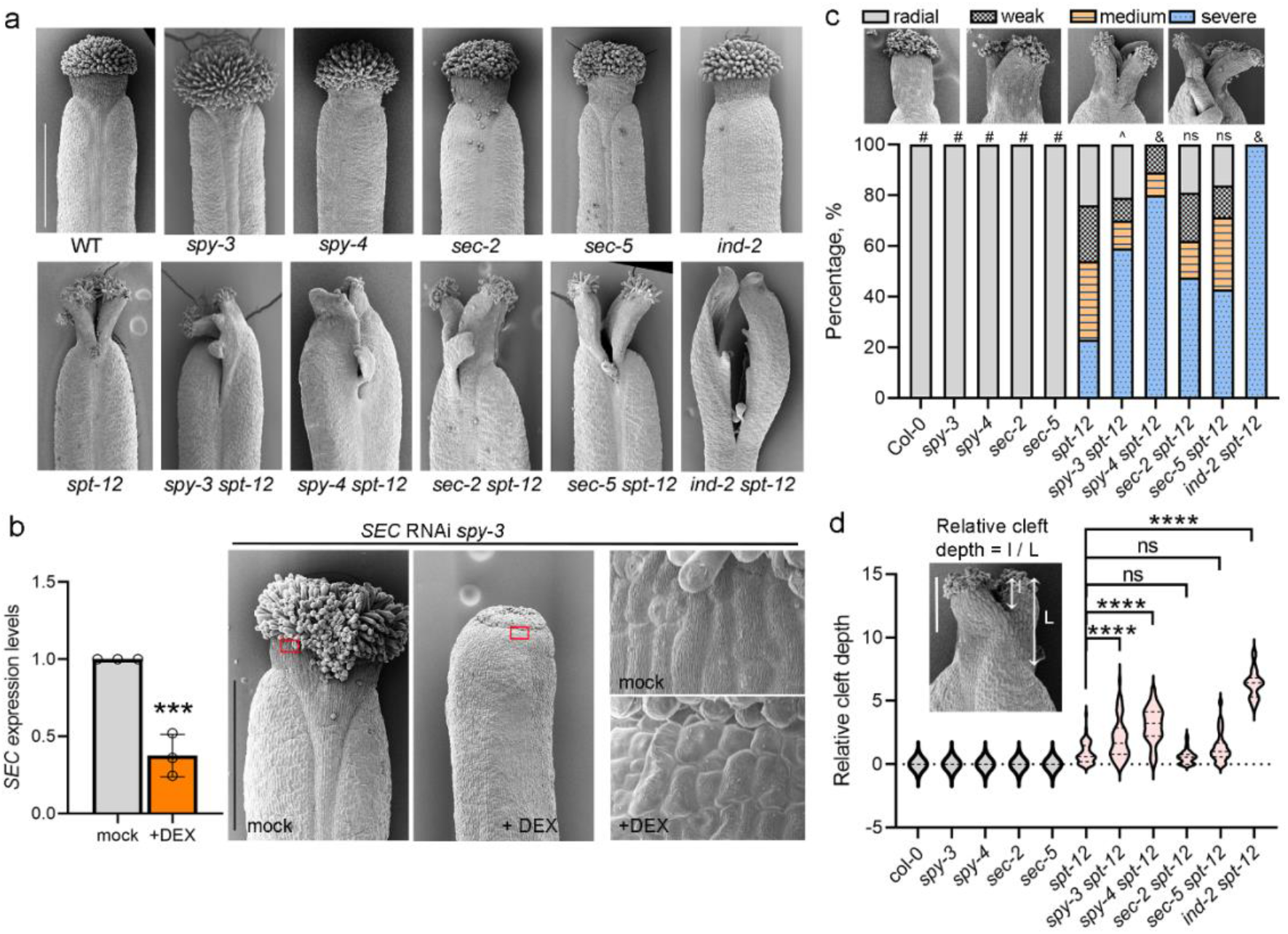
Loss of SEC and SPY function leads to defects in style development and enhances symmetry breaking in the *spt-12* mutant style. **a**, Representative SEM micrographs of stage12 gynoecia from the genetic backgrounds depicted on the panels. Scale bar indicates 500 μm. **b**, Analysis of *SEC* RNAi *spy-3* transgenic line showing downregulation of *SEC* expression levels after DEX-treated inflorescences compared to mock treatments, by qRT-PCR (left panel) and SEM analysis of mock- and DEX-treated gynoecia (right panel) showing defects in style development, *i.e.*, strong reduction of style development/growth. Micrographs on the far-right-hand side show close-up images of the style regions (red boxes) displaying morphological cell defects. Scale bar indicates 500 μm. **c**, Quantification of the frequency and severity of the bilaterally symmetric defective styles observed in the double *sec spt* and *spy spt* mutants compared to their parental lines. Phenotypes were grouped in the following four categories: Radial (WT-like style, radially symmetric); weak (shallow cleft at the organ distal tip); medium (cleft running through the middle of the style); severe (deep cleft spanning the style into the ovary region). Fifty gynoecia were analysed for each genotype. Phenotypic classes were compared with *spt-12* using 4 × 2 contingency tables followed by Pearson’s χ^2^ test. Two-tailed P values are as follows: Col-0 vs *spt-12*, P< 0.00001 (#); *spy-3* vs *spt-12*, P< 0.00001 (#);*spy-4* vs *spt-12*, P< 0.00001 (#); *sec-2* vs *spt-12*, P< 0.00001 (#); *sec-5* vs *spt-12*, P< 0.00001 (#); *sec-2* vs *spt-12*, P< 0.00001 (#);*spy-3 spt12* vs *spt-12*, P = 0.000019 (^); *spy-3 spt12* vs *spt-12*, P = 0.000019 (^); *spy-4 spt12* vs *spt-12*, P< 0.00001 (&); *sec-2 spt-12* vs *spt-12*, P = 0.0249 (ns); *sec-5 spt-12* vs *spt-12*, P = 0.1007 (ns); *ind-2 spt-12* vs *spt-12*, P < 0.00001(&). *P* values < 0.01 were considered as extremely statistically significant. **d**, Quantification of the severity of the bilateral split style (Relative cleft depth) was measured as the depth of the cleft (I) relative to the style length (L). Fifty gynoecia were analysed for each genotype. Phenotypic classes were compared with *spt-12* using one-way ANOVA multiple comparisons: ****, P < 0.0001; ns, no significant, P > 0.05.

This suggests a synergistic and/or redundant role for both enzymes in controlling the morphogenesis of the gynoecium apex. To overcome the embryo lethal effect elicited by the elimination of both enzymatic activities^32,37^ and shed light on the synergistic post-embryonic roles of the two enzymes-specifically during style development- we produced a dexamethasone (DEX) inducible SEC *RNAi* construct to lower *SEC* mRNA levels in the *spy- 3* background (*spy-3,SEC RNAi*) (Extended Data Fig. 5a). Three independent *Arabidopsis* transgenic lines, which displayed reduced levels of *SEC* transcripts by qRT-PCR experiments (Extended Data Fig. 5b) were further analysed by SEM for phenotypical analysis. We observed defects in style formation after DEX treatment *i.e*., strong reduction in style length, lack of wax/crenulations of stylar cells (used as a differentiation marker for style) (Fig. 3b) and a consistent reduction in fruit length (Extended Data Fig. 5c). These results revealed a previously unrecognised role for SEC and SPY in style (and fruit) development and suggest a synergistic, additive control of these *O*-glycosyltransferases on style-specific targets.

To investigate the role of SEC and SPY in style radial symmetry establishment via regulation of SPT activity, we analysed the gynoecia of *spt sec* (*spt-12 sec-5* and *spt-12 sec-2*) and *spt spy* (*spt-12 spy-4* and *spt-12 spy-3*) double mutants (Fig. 3a). Our genetic analysis showed that *spt sec* double mutants displayed a similar phenotype as compared to the segregating *spt-12* control, while *spt spy* double mutant combinations increased the frequency and severity of the *spt* split-style phenotype, by strongly augmenting the percentage of bilateral vs radial styles observed (Fig. 3c), as well as the depth of the medial cleft (measured as the ratio between the length of the cleft vs the style length) (Fig. 3d).

These data indicate that *spt* is epistatic to loss of both SEC and SPY enzymatic activities and suggests that other key regulators of style development may be regulated by SPY. To further examine this, *sec* and *spy* mutant gynoecia were treated with CK. No defects were displayed by *sec* mutants, however for the *spy* mutants we observed extensive proliferation of the medial-apical region that connects stigma and replum (*spy-3*) as well as ectopic, unbalanced growth of the lateral shoulders (*spy-4*) (Extended Data Fig. 6), a novel phenotype associated with CK hypersensitivity which has not been previously observed. These data not only provide the first genetic framework for a SPY-CK antagonistic interaction, but also corroborate the idea that SPY and SPT work on similar pathways, as they both repress the cytokinin proliferating output at the gynoecium apex.

### Specific Ser/Thr residues at the N-terminus of SPT promote radial style development

To test the effect of *O*-glycosyl decorations on SPT function during radial symmetry establishment at the gynoecium apex we carried out a detailed mutant complementation assay by producing a series of loss-of-*O*-glycosylation mutant variants of SPT (expressed under the SPT native 5-Kb promoter^39^) and analysed their ability to complement the *spt* split-style phenotype. To this end, specific Ser (S) and Thr (T) residues targeted by SEC and SPY *in vivo* (Fig. 1a) and *in vitro* (Extended Data Fig. 3b), were mutated to alanine (Ala, A) to mimic loss of modification^40^. We predicted the lack of style complementation would indicate a positive functional role for those residues and the associated PTMs in sustaining SPT function. We produced and analysed the following mutant versions of SPT: S23-to-A; S23, S24; S25-to-A (hereafter S23-25-to-A); S60-S61-to-A; T71, T72, T73, T74-to-A (hereafter T71-74-to-A); S23, S24, S25, T71, T72, T73, T74-to-A (hereafter S+T-to-A) (Fig. 4a). Firstly, to exclude the possibility that changes in the amino acid sequence of SPT would lead to alterations in transcript levels and protein stability or localization, we fused the SPT wild-type and mutant sequences with yellow fluorescent protein (YFP) and carried out qRT-PCR experiments (Extended Data Fig. 7) and confocal microscopy analysis (Fig. 4b). None of the lines examined showed a significant reduction in *SPT* mRNA levels (Extended Data Fig. 7). Furthermore, confocal microscopy analysis showed clear YFP signals present at the apex of gynoecia expressing the wild-type SPT complementation line as well as the S23-to-A (peptide_1) and S60,S61-to-A (peptide_2) point mutations (Fig. 4b). Accordingly, SEM analysis showed that the *spt* phenotype was complemented by S23-to-A and S60,S61-to-A mutations to a similar level as the wild-type sequence (Fig. 4b). On the other hand, despite clear SPT nuclear expression at the gynoecium apex of S23-25-to-A (peptide_1), T71-74-to-A (peptide_2) and S+T-to-A (both peptides), these point-mutations lines displayed bilateral styles, similar to the *spt* mutant (Fig. 4b).

**Fig. 4.**
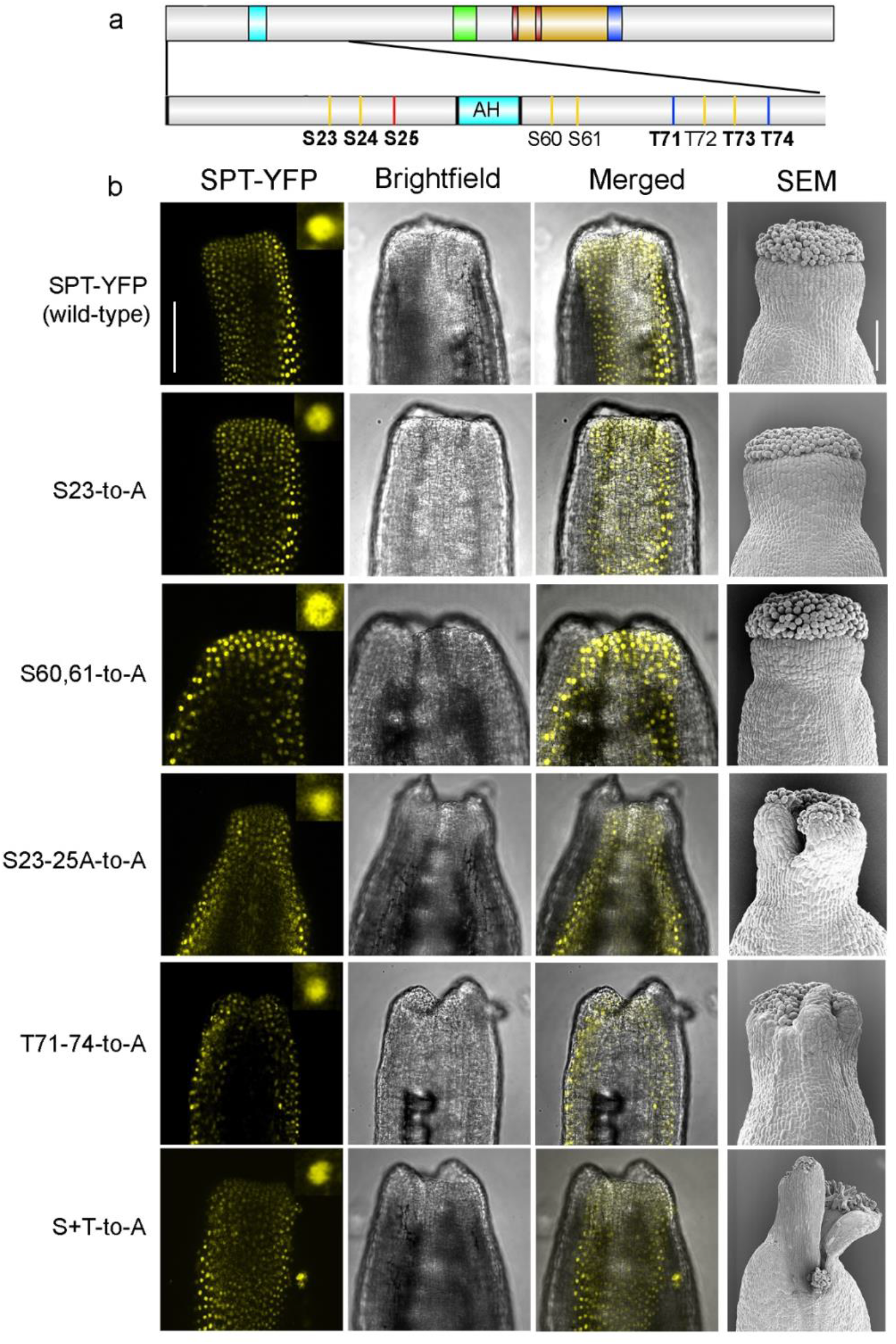
Genetic complementation analysis of SPT revealed positive role for *O-*glycosyl modified residues in radial style development. **a**, Schematic representation of SPT residues modified *in vivo* and *in vitro* by *O*-fucose (red and yellow) and *O*-GlcNAc (blue and yellow) for point mutations: Ser (S) and Thr (T) residues depicted in the panel were mutated to Ala (A). **b**, Representative confocal and SEM images of gynoecia of the SPT-YFP complementation and point-mutation lines, showing YFP signal at the style and their respective style phenotype, radial vs bilateral style. S23-25-to-A (S23,S24,S25-to-A); T71-74-to-A (T71,T72,T73,T74-to-A); S+T-to-A (S23,S24,S25,T71,T72,T73,T74-to-A). Scale bars indicate 500 μm. Inset: enlarged nucleus with SPT-YFP signal from the apex of gynoecia.

Thus, these results provide a functional role for specific SPT residues in symmetry establishment and suggest that the PTMs attached to these residues are fundamental for SPT function in style development via a novel mechanism distinct from transcriptional expression, protein stability and cellular localisation.

### *O*-glycosyl PTMs on SPT promote its transcriptional activity

Since SPT binds DNA – and thus regulates transcription - as a dimer, we hypothesised SEC and SPY may impact its ability to interact with protein partners and/or specific promoters. To shed light on how these PTMs of SPT may play a role in orchestrating radial style formation, we tested whether SEC and SPY could enhance the formation of the SPT-SPT homodimers and/or SPT-IND and SPT-HEC1 heterodimers essential for style development. We performed FRET-FLIM quantitative assays in tobacco leaves to assess whether the co-expression of agrobacteria harbouring either SEC:HA (*p35S:SEC-HA*) or SPY:HA (*p35S:SPY-HA*) would lead to a further reduction in the lifetime of the FRET donor (GFP) as compared to the formation of the homodimer SPT-GFP;SPT-RFP (*p35S:SPT-GFP*;*p35S:SPT-RFP*), and the heterodimers SPT-GFP; IND-RFP *(p35S:SPT-GFP*;*p35S:IND-RFP*) and SPT-GFP; HEC1-RFP (*p35S:SPT-GFP*;*p35S:HEC1-RFP*).

To begin, we confirmed the expression of SEC-HA and SPY-HA recombinant proteins by western blotting (Extended Data Fig. 8a) and direct interactions in the nuclei between SPT-GFP;SPT-RFP as well as SPT-GFP;IND-RFP and SPT-GFP;HEC1-RFP (without co-expression of the enzymes) with FRET-FLIM assays. A significant reduction of the GFP fluorescence lifetime in the presence of SPT-GFP;SPT-RFP, SPT-GFP;IND-RFP and SPT-GFP;HEC1-RFP interactions was detected as compared with the fluorescence lifetime of the GFP in the negative control: SPT-GFP;RFP-NLS (which targets the RFP protein in the nucleus) (Fig. 5a). These changes in GFP fluorescence lifetime are consistent with FRET efficiencies of 5.7 %, 9.6 %, and 7.4 % for SPT;SPT, SPT;IND, and SPT;HEC1, respectively (Fig. 5a). The positive FRET control, SPT-GFP-RFP (in which the two florescent tags are both cloned in cis to SPT) showed an even stronger decrease in GFP fluorescence lifetime, which translated into a FRET efficiency of 18.8 % (Fig. 5a). Thus, the presence of either SEC-HA or SPY-HA had no significant effect on the FRET efficiencies observed in the presence of interactions between SPT-GFP and its RFP-tagged partners (Fig. 5a). In principle, this would suggest that *O*-GlcNAc and *O*-fucose do not promote the formation of SPT-containing dimers. However, the enzymes may modulate dimer formation via mechanisms that are incompatible with a FRET increase, e.g., by conformational rearrangements that increase the distance between the FRET donor and acceptor.

**Fig. 5.**
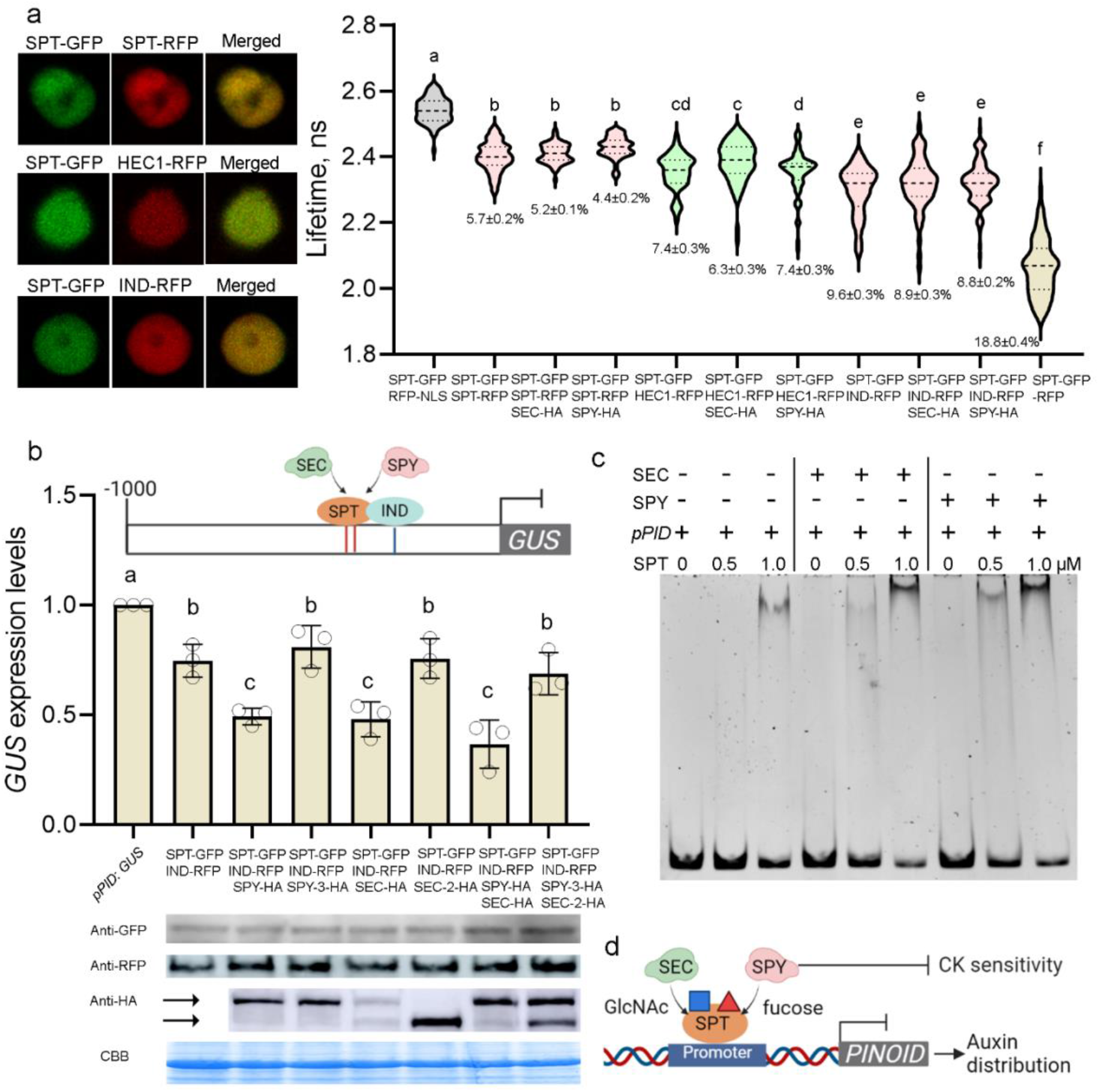
SEC and SPY promote the SPT-mediated repression of *PID* transcription rather than its ability to form homo- and hetero-dimers. **a**, FRET-FLIM assay showing both SEC and SPY have no effect on the strength of the interaction between SPT-GFP and its interacting partners SPT-RFP, IND-RFP and HEC1-RFP, in nuclei (left panels) of co-infiltrated tobacco leaves. The violin plots represent the distribution of lifetime values, the dotted lines within violin represent the median (central line), upper and lower quartiles. Experiments were repeated at least three times for each combination; more than 60 nuclei were used for quantification for each combination. The FRET efficiency (%) values (means±SE) are also indicated in the graph. Different letters indicate significant difference (*p*< 0.01) tested by one-way ANOVA multiple comparisons. **b**, Schematic representation of the 1-Kb promoter of *PID* used in transactivation assays, including the TF binding sites (G-boxes, red lines, and E-box, yellow line). Quantification of *GUS* expression by qRT-PCR experiments showing both SEC-HA and SPY-HA enhance the transcriptional downregulation triggered by SPT and IND. Here the OD values used for both SPT and IND were 0.1 in each combination. Experiments were repeated at least three times for each combination. Values shown are means±SD. Different letters indicate significant difference (p<0.0001) tested by one-way ANOVA multiple comparisons. Bottom: immunodetection of SPT-GFP, IND-RFP, SEC-HA, SEC-2-HA, SPY-HA and SPY-3-HA from *N. benthamiana* Agro-infiltrated leaves used for transactivation assays. The upper arrow indicates SEC-HA, SPY-HA or SPY-3-HA while the lower arrow indicates SEC-2-HA. The CBB (Coomassie Brilliant Blue) band was used as sample loading control. **c**, EMSA experiments showing SEC (5TPR-SEC) and SPY (3TPR-SPY) both enhance binding of SPT (6xHis-SPT) to the region of *PID* promoter (171-bp p*PID*). Similar results were obtained from two independent experiments. **d**, Schematic working model showing SEC and SPY act upstream of SPT to attach *O-*GlcNAc (red square) and *O-*fucose (blue triangle), which in turn promotes binding of SPT to the *PINOID* promoter and/or its transcriptional activity. This way, SEC and SPY contribute to the fine-tuning of auxin distribution while repressing CK sensitivity at the gynoecium apex, ultimately promoting organ symmetry transition.

Next, we investigated whether SEC and SPY would impact SPT-mediated gene expression. The best characterised downstream target of SPT (and IND) during radial symmetry establishment is the *PINOID* (*PID*) kinase^8,12,21,41^ which is downregulated to support the accumulation of the auxin morphogenic signal within specific medial-apical cells [8]. Thus, we tested whether downregulation of *PID* expression by SPT-IND was exacerbated by SEC and SPY by performing a transient trans-activation assay in *N. benthamiana* leaves using the same constructs used for the FRET-FLIM experiments and a 1-Kb promoter fragment of *PID*, containing the *cis-*elements directly bound by both IND^21,41^ and SPT *in vivo* (Extended Data Fig. 8b).

The *PID* promoter was fused to the *GUS* gene as a reporter. To begin, we confirmed the expression of SEC-HA or SPY-HA didn’t affect *GUS* expression (Extended Data Fig. 8c), and the transcriptional readout of SPT/IND was determined by qRT-PCR experiments using different OD for co-infiltration (Fig. 5b and Extended Data Fig. 8d). Firstly, we tested for an equal expression of the co-infiltrated recombinant proteins by western blots (Fig. 5b) and then compared the *GUS* transcription levels. As expected, expression of *SPT-GFP* and *IND-RFP* either alone (homodimers) or co-expressed (heterodimer) significantly diminished *pPID:GUS* expression levels (Fig. 5b and Extended Data Fig. 8d), while expression of either *SEC-HA* or *SPY-HA* had no significant effect on transcription alone (Extended Data Fig. 8c). On the other hand, co-expression of both *SEC-HA* and *SPY-HA* together with *SPT-GFP* and *IND-RFP* led to a significant and synergistic further reduction of the background levels of *PID* expression (Fig. 5b). Furthermore, co-expression of the catalytic *SPY* (*SPY-3-HA*) and *SEC* (*SEC-5-HA*) mutant enzymes together with SPT-GFP and IND-RFP recombinant proteins, led to a reversion of the *GUS* transcriptional levels to the basal repression trigged by the SPT-GFP/IND-RFP heterodimer (Fig. 5b). Altogether, these results demonstrate that both SEC and SPY enhance SPT/IND-mediated repression of *PID* transcription.

To corroborate the idea that *O*-glycosyl PTMs of SPT directly influence its transcriptional activity rather than dimerization events, we tested whether the affinity of SPT for the *PID* promoter was enhanced by SEC and SPY.

To this end, we employed EMSA experiments performed using the full-length 6xHis-SPT recombinant protein (Extended Data Fig. 3a) which we demonstrated was able to bind a 171-bp fragment of the *PID* promoter encompassing the *cis*-elements (G-box, CACGTG) (Extended Data Fig. 8e) recognised *in vivo* by SPT (Extended Data Fig. 8b). Moreover, the electrophoresis mobility shift signal was strongly abolished when the wild-type G-box sequence of the *PID* promoter was mutated (Mut, TGATGA)^42^ (Extended Data Fig. 8e) supporting previous data^21^ which demonstrated SPT binds to this specific region of the *PID* promoter both *in vivo* and *in vitro*. Notably, when SPT was incubated with the *PID* promoter fragment in the presence of recombinant 5TPR-SEC-HA and 3TPR-SPY-HA proteins and their respective donor substrates, we observed a stronger mobility shift occurring at lower concentrations of SPT (Fig. 5c).

Altogether, our findings show that SEC and SPY can modulate SPT-mediated control of *PID* expression by increasing its DNA-binding affinity and/or transcriptional activity rather than its ability to form homo- or hetero-dimers.

## DISCUSSION

This work provides the genetic and molecular basis for the role of *Arabidopsis* SEC^29^ and SPY^30^ *O*-glycosyl enzymes in the regulation of SPT, a bHLH TF critical for radial style development^8,18^. More broadly, our data provide the first molecular link between *O*-fucose and *O*-GlcNAc modifications in symmetry foundation during multicellular development, by unveiling the upstream control of a key regulator of radial style development of *Arabidopsis thaliana*. As comparable biological processes and conserved molecular players *i.e*., *O*-GlcNAc^26^ and bHLH TFs^43^ are present in both plant and animal kingdoms, this study sheds light on a fundamental mechanism which could be employed more generally during the orchestration of symmetry establishment in high eukaryotes.

Our genetic analysis and inducible *sec spy RNAi* mutant (Fig. 3) unveiled fundamental roles for these two enzymes in style development and show how the synergistic activity of *O*-GlcNAc and *O*-fucose contribute to the foundation of plant organ symmetry.

Accordingly, the importance of *O*-GlcNAc PTM in metazoan development is well known due to the extreme phenotypes and embryo lethality displayed by loss-of-function mutants for the *O*-GlcNAc transferase (OGT) enzyme^26,27,44^. During style development, the *O*-GlcNAc/*O*-fucose acceptor-substrate investigated in this work, SPT, plays pivotal roles to specify the medial tissue identity^18^, coordinate the medio-lateral^19^ and adaxial-abaxial^9^ body axes and control the antagonistic auxin/cytokinin balance, to establish radial symmetry^8,16,45^.

Here we proved that SPY and SEC both physically interact with SPT *in vivo* and *in vitro* (Fig. 2) and modify specific Ser/Thr residues at the N-terminus of SPT (Fig. 1) flanking its AH domain which is also sufficient for interactions with the enzymes (Fig. 2). It has long been known that the AH domain of SPT supports the essential role of the Acidic domain in carpel function^36^, but the molecular basis of its modulation is yet to be determined. Since, the N-terminal region, including both AH and Ac domains is important for SPT transcriptional activation^36^, the attachment of PTMs flanking the AH domain support two mechanistic scenarios for the function of the AH helix as a modulator of the Ac domain activity: (i) a direct effect via a conformational change or (ii) an indirect effect via interaction with other transcriptional activators and/or co-repressors.

Moreover, the AH domain of SPT is quite diverse from the N-terminus sequence of other plant bHLH TFs (except for the SPT closest homolog, ALCATRAZ)^36^ reinforcing the idea that it hosts specific residues for signal transduction mediated by *O*-GlcNAc and *O*-fucose. Notably, the AH domain has been found either highly conserved across some SPT orthologues (*e.g*., tomato) or missing from other plant species completely – e.g., in monocot rice^36^. This supports a scenario in which evolution of the N-terminus of SPT and its regulation by sugar decorations might be linked to the diversification of gynoecia style/stigma shapes, *i.e*., bilateral in monocots and radial in dicots.

*O*-GlcNAc has been linked to the cold-temperature response in both animals^46^ and plants^47^. Interestingly, ambient temperature correlates with cellular levels of *O*-GlcNAc PTMs, a signalling mechanism frequently employed to confer protection against stresses linked to high and low temperatures in animal systems^46^. Accordingly, reduced *O*-GlcNAc levels in hypomorphic *ogt* mutants in *Drosophila* lead to increased larval/pupal lethality at higher temperatures^48^. Interestingly, in addition to being a master regulator of organ symmetry establishment, SPT can transduce environmental cues into developmental programmes, by integrating external signals such as cold temperature^49,50^ and light quality^51^ during seed germination and style development respectively. Even though SPT was the first regulatory gene described to control the cold response during seed germination^49^, the molecular mechanism underpinning this process is still unelucidated, and post-translational control of SPT was hypothesised to underpin its mechanistic activity^50^. Thus, this raises the intriguing possibility that O-GlcNAcylation might regulate SPT at the post-transcriptional level to integrate environmental cues into organ development.

A recent *O*-fucosylomics^52^ study performed in young *Arabidopsis* seedlings, identified a N-terminal peptide of SPT modified by *O-*fucose, coinciding with our peptide_1 (Fig.1), whose modification was lost in *spy-4* mutant background. Thus, this strengthens the importance of sugar-based post-translational modification of SPT in other developmental contexts and such regulatory mechanism would fit well with the view of SPT as integrator of abiotic signals and genetic factors, similar to that proposed for the DELLA protein RGA^53^.

It has been previously proposed that SEC and SPY control the activity of other plant acceptor substrates by promoting protein stability^54^, cellular localization^55^, enhancing or disrupting protein–protein interactions^30,56^ and orchestrating gene expression^55^. Amongst these, the best characterized molecular example is the DELLA protein RGA, which is antagonistically regulated by SEC and SPY during *Arabidopsis* post-embryonic development^29,30^. Interestingly, the synergistic activity of SEC and SPY on SPT represents, to the best of our knowledge, the first example of both *O*-GlcNAc and *O*-fucose modification synergistically regulating a target protein during post-embryonic plant development.

The complex network of transcription factors orchestrating style development includes several layers of cross- and self-regulation, that act at both the transcriptional and translational levels^17^. The importance of dimerisation events for bHLH TF activity at the gynoecium apex and the spatio-temporal activity of these master regulators highlights how the post-translational regulation by *O*-GlcNAc and *O*-fucose may act in a cell-/tissue-type manner and/or at a specific developmental stage. Although our FRET-FLIM experiments in tobacco leaves indicate SEC and SPY may not regulate SPT-SPT, SPT-IND, and SPT-HEC1 dimer formation (Fig. 5), it is still possible that a gradient of TF interactions at a protein level might be mediated by *O*-GlcNAc and *O*-fucose specifically at apical-medial gynoecium cells undergoing symmetry transition. Also, additional upstream regulation can be hypothesised, as three NGA family members have been identified as targets of *O*-GlcNAc modification in *Arabidopsis* inflorescences^25^. This scenario is also compatible with the prediction that other key regulators of style development are *O*-fucosylated by SPY, which would explain the additive, synergistic phenotype observed in the *spt spy* mutant resembling *spt ind^21^* (Fig. 3a) and *spt hecs* phenotypes^9,16^.

Taking the data as a whole, we propose a working model (Figure 5d) in which *O-*glycosylation of SPT by *O*-GlcNAc and *O*-fucose is mediated by SEC and SPY, respectively, and regulates style radial symmetry establishment at the gynoecium apex by enhancing the SPT-mediated repression of *PID* transcription, ultimately balancing the auxin/cytokinin crosstalk required for style development.

## Acknowledgments

We are much obliged to Prof. Neil E Olszewski (University of Minnesota, USA) for providing the *spy-3, spy-4 and sec-2* mutants, Prof. Lijing Xing (Institute of Botany, CAS, China) for the *sec-5* mutant line and Prof. Dolf Weijers (Wageningen University, The Netherlands) for the SPT translational fusion line *pSPT:SPT-sYFP*. We are thankful to Dr. Yuli Ding, Dr. Shujuan Xu and Dr. Estee Tee (JIC) for help in setting up Co-IP & split-Luciferase complementation assays, ChIP assay, and FRET-FLIM assay respectively, and Prof. Lei Wang (Institute of Botany, CAS, China) for useful suggestions for setting up the *in vitro* modification assay. We also thank Dr. Lipeng Feng and Prof. Tony Maxwell (JIC) for helping to setup the EMSA assay. We are grateful for the access to the JIC Bioimaging Facility including staff assistance and training, as well as the Horticultural Platform. We thank Prof. Tai-ping Sun (Duke University, North Caroline, USA) for discussions and helpful comments on the manuscript.

## Author contributions

L.M. conceptualized the project; Y.J. and L.M. designed the experiments; Y.J. performed most of the experimental work with help from S.C-F, S.K., I.J., L.A., S.L., C.M., G.S. and L.M.; All authors analysed the data; Y.J. prepared the figures; L.M. wrote the manuscript and all authors commented and edited the manuscript.

## Author Information

The authors declare no competing financial interests. Readers are welcome to comment on the online version of the paper.

## Data and materials availability

All data needed to evaluate the conclusions in this paper are present in the paper and/or the Supplementary Materials. All raw proteomic data are available on PRIDE repository (https://www.ebi.ac.uk/pride/, accession number: PXD037917). All new genetic material (high-order mutants, transgenic lines) and expression vectors will be made available to the scientific community upon request and with no limitation.

## Methods

### Plant materials and growth conditions

The following loss-of-function mutant lines in ecotype Columbia (Col-0) background were used in this study: *sec-2^32^*, *sec-5*^34^, *spy-3^35^*, *ind-2^24^*, *spt-12^57^*. The *spy-4* mutant^35^, which was originally in Wassilewskija background, was back-crossed to Col-0 three consecutive times and homozygous segregating seeds were used for this study. The following transgenic line was described previously: *pSPT:SPT-sYFP^39^*. The *pSPT:SPT-sYFP* in Col-0 background was crossed with *spt-12* to obtain the complementation line used in this study. Further, the *pSPT:SPT-sYFP spt-12* line was crossed with *sec-5* and *spy-3* mutants. Plants were grown at 22 °C in long-day conditions (16 h light / 8 h dark) in controlled environment rooms.

### DNA constructs

DEX inducible amiRNA for *SEC* was constructed as following: the primers for amiRNA generation was chosen as previously shown^58^. Using *pRS300^58^* as template, the amplified fragment was inserted downstream of a Dexamethasone-inducible promoter to generate the *pGAL6:amiRNA* plasmids. The resultant plasmids were recombined with *p35S:GVG* plasmid and the Basta selection marker cassette (Synbio TSL) using standard Golden-Gate cloning methods to produce the binary vector *p35S:GVG-amiSEC-Basta-pICSL4723 (p35S: amiSEC-GR)*, which was transformed into the *spy-3* mutant background.

To generate the transgenic SPT-sYFP point mutation lines analysed in this study, we amplified a 5-kb *SPT* promoter fragment from Col-0 genomic DNA using the primers pair (see Supplementary Table 1 for sequence): pSPT-F and pSPT-R. The *SPT* promoter was cloned into *pCambia1305* vector by using *EcoRI* and *BspEI* restriction enzymes, thus producing *pSPT-pCambia1305* plasmid. Then the *SPT* wild-type genomic coding sequence fused in frame to the sYFP fluorescent tag (*SPT-sYFP*) was amplified as a whole segment (2.5 kb) from genomic DNA extracted from *pSPT:SPT-sYFP^39^* seedlings using the primers pair 2.5k_(BspEI)_F and 2.5k_(KpnI)_R and cloned into the *pSPT-pCambia1305* plasmid, thus obtaining the *pSPT:SPT-sYFP-pCambia1305* construct.

To generate constructs harbouring specific point mutations of SPT, we used the *pSPT:SPT-sYFP-pCambia1305* plasmid as template and used mutagenesis primers (listed in Supplementary Table 1) introducing specific mutations in the SPT coding sequence. All constructs were verified by sequencing and introduced into *Agrobacterium tumefaciens GV3101* strain by heat shock, then transformed into *spt-12* background by floral dipping (note, since *spt-12* produces few seeds, each construct was transformed in 30 mutant plants). The transgenic seeds were selected on MS plates supplied with 15 mg/litre Basta and at least eight positive, independent T1 lines were screened at the confocal microscope for the presence of nuclear YFP signals in roots.

To produce the recombinant 6xHis-SPT protein, the full-length *SPT* coding sequence was cloned into *pRSFDuet* vector by using *BamHI* and *PstI* restriction sites. To produce recombinant 5TPR-MBP-SEC and 3TPR-MBP-SPY proteins, we use the following strategy: Firstly, to generate a protein expression vector with 10xHis-MBP tag, we amplified a 10xHis-MBP coding fragment by PCR with primers pair 10xHis-MBP-F and 10xHis-MBP-R from *pMAL-c2X*® vector (Addgene), and cloned it into *pTrcHis*® vector (Addgene) using *XhoI* and *HindIII* restriction enzymes, obtaining a *pTrc10xHis-MBP* vector. Secondly, code-optimised full-length *SEC* and *SPY* coding sequences were synthesized by Sangon Biotech (Shanghai). The coding sequences of *5TPR-SEC* and *3TPR-SPY* were amplified using gene-specific primers pairs 5TPR-SEC-opt_F/5TPR-SEC-opt_R and 3TPR-SPY-opt_F/3TPR-SPY-opt_R and cloned into the *pTrc10xHis-MBP* vector using *SacI* and *SalI* restriction enzymes. All constructs were verified by sequencing and transformed into *E. coli* Rosetta (DE3) strain by heat shock.

For transient co-expression in tobacco, *35S::SPT-3xFLAG*, *35S::SEC-3xHA* and *35S::SPY-3xHA* were constructed as follows: the full-length coding sequences of *SPT, SEC*, and *SPY* were amplified using gene-specific primers (listed in Supplementary Table 1), the PCR products were digested with *SifI* (Takara) and cloned into the empty *pCambia1305-3xFLAG* or *pCambia1305-3xHA* vectors (gifted by Dr Yuli Ding) which were pre-digested with *DraIII* (Takara). All constructs were verified by sequencing and introduced into *Agrobacterium tumefaciens GV3101* strain for transformation in *N.benthamiana* leaves.

For split-Luciferase complementation (Split-Luc) assay, the coding sequence of *SPT, SEC* and *SPY* were amplified by PCR using gene-specific primers (listed in Supplementary Table 1) and inserted into *pCambia1305-35S::Nluc* and *pCambia1305-35S::Cluc* vectors (gifted by Dr Yuli Ding), respectively, using *DraIII* restriction sites, thus producing the *35S::SPT-Nluc*, *35S::SEC-Cluc* and *35S::SPY-Cluc* constructs.

For the FRET-FLIM assay, firstly the coding sequence of *SPT, HEC1* and *IND* were inserted into *pCambia1305* using *DraIII* sites, then the coding sequences for *EGFP* or *RFP* were inserted in frame using *XbaI* and *PstI* restriction enzymes.

For the transactivation assay, we firstly cloned the *GUS* coding sequence into the *pCambia1305* vector using *HindIII* and *BstEII* sites, producing the *pCambia1305-GUS* contruct. Then the 1-kb promoter region of *PINOID* was cloned in *pCambia1305-GUS* vector using *PstI* and *HindIII* restriction sites. All constructs were verified by sequencing and introduced into *Agrobacterium tumefaciens GV3101* strain for transformation in *N.benthamiana* leaves.

For Y2H assay, full-length and domain fragments of coding sequences from SPT, SEC and SPY were cloned into *pDONR207*® (Invitrogen) using primers in Supplementary Table 1, and then recombined into either *pGDAT7*® or *pGBKT7*® vectors (Clontech) following the manufacture instructions.

For EMSA assay, the 171 bp fragment of *PID* promoter was amplified from *Arabidopsis* genomic DNA with primers (listed in Supplementary Table 1). To test the specificity of SPT binding to the DNA fragment, a 171 bp DNA fragment with mutated G-boxes was synthesized (Sigma).

### Gynoecium treatment

For dexamethasone (DEX) treatments, inflorescences from the *p35S: amiSEC-GR* inflorescences were sprayed with either 10 µM DEX (Sigma) or mock (DMSO, Sigma) three times, every 5 days over two weeks, and gynoecia were fixed after 5 days from the last treatment. For the cytokinin (BA) treatments, the inflorescences of Col-0, *sec-2*, *sec-5*, *spy-3* and *spy-4* were sprayed with either 50 µM or 100 µM BA (6-benzylaminoadenine, Sigma B3408) or mock [200 mM NaOH] three times, every 5 days over two weeks, and gynoecia were fixed after 5 days from the last treatment. All spray treatments used 0.015% final concentration of Silwet L-77.

### Scanning electron microscopy

Inflorescences were fixed overnight in FAA solution (3.7% formaldehyde, 5% glacial acetic acid and 50% ethanol). After complete dehydration through an ethanol series from 50% until 100%, the inflorescences were critical point dried using the Leica EM CPD300. Gynoecia were hand-dissected using a stereomicroscope (Leica S9D), coated with gold particles and examined with FEI Nova NanoSEM 450 emission scanning electron microscope using an acceleration voltage of 3 kV. Experiments were conducted in biological triplicates and gynoecia were taken from distinct inflorescences and plants each time. The total number of gynoecia analysed for each experiment is indicated in the figure legends.

### RNA extraction and qRT-PCR

Three independent biological repeats of total RNA extracted from each genotype were isolated from young inflorescences using RNeasy Plant Mini Kit (Qiagen), including treatment with RNase-free DNase (Qiagen) following the manufacturer’s instructions. Four microgram of extracted RNA sample was reverse transcribed using the M-MLV Reverse Transcriptase (Promega) from each RNA sample. The qRT-PCR experiments were performed in quadruplicates using SYGREEN BLUE qPCR MIX (PCRBIO) with Chromo4 Real-Time PCR Detection System (Bio-Rad). Target gene expression level was quantified by the 2^−ΔΔct^ method with *UBIQUITIN10* as internal control. Gene-specific primers are listed in Supplementary Table 1.

### Confocal microscopy

Confocal microscopy analysis was performed on a Zeiss LSM 880 confocal microscope with a 40x water immersion lens, using the ZEN-Black-LSM880 software (Zeiss, Germany). YFP signals were excited by an Argon-514 nm, 10 mW solid-state laser with emission at 550-570 nm. To view the gynoecia, floral buds were dissected using a stereomicroscope (Leica S9D), then the gynoecia were mounted along their longitudinal axis in water. To quantify the YFP fluorescence intensity, the z-series images of epidermal cells on the surface layer (L1) at the style region were collected with the z-step set at 1μm. Maximum intensity projection of z-series of nucleus was used to quantify the YFP intensity using the software ImageJ (v1.53).

### Immuno-precipitation from Arabidopsis inflorescences

*SPT-sYFP* was purified from young inflorescences of *SPT-sYFP spt-12* line (segregated control), *SPT-sYFP spt-12 sec-5* and *SPT-sYFP spt-12 spy-3* plants. Young buds (close sepals) were collected from the inflorescences and immediately frozen in liquid N_2_ after picking, five grams were used in each of the three biological replicates for each genotype. Each sample was grinded in liquid N_2_ and extracted using 10 ml buffer A as previously described ^55^. After centrifugation at 14,000 rpm for 30 min, the supernatant was incubated with 30 μl GFP-Trap beads (ChromTek). After 1 h rotation at 4 °C, the beads were sedimented with a magnetic rack (GE Healthcare) for 1 minute and washed with buffer A for three times. Proteins on beads were eluted with 1x loading buffer (sigma), then separated by 10% SDS PAGE and used for Mass Spectrometry analysis.

### Recombinant proteins expression

Cells were grown in LB media at 37°C until OD600 value reached 0.5, then the culture was cooled down before induction. The expression of 6xHis-SPT was induced by 0.4 mM IPTG at 25°C for 4 h. The expression of *10xHis-MBP-5TPR-SEC* and *10xHis-MBP-3TPR-SPY* were induced by 0.4 mM IPTG at 16°C for 16 h. Recombinant proteins were purified with Nickel sepharose according to the manufacturer’s instructions (QIAGEN). The purified recombinant protein was dialyzed overnight against dialysis buffer (20 mM Tris-HCl, pH 8.0, 50 mM NaCl, 1 mM DTT, 5% glycerol) at 4°C. All proteins were aliquoted, flash-frozen in liquid N_2_ and stored at -80°C.

### *In vitro O-*glycosylation assay

The direct modification of SPT protein by SEC and SPY were tested by an *in vitro* glycosylation assay as previously described in Zentella et al^30^. Briefly, to test *O*-GlcNAcylation, a 50 μl reaction was carried out by mixing 10 μg *SPT-FLAG*, 5 μg *5TPR-SEC*, 20 mM Tris-HCl, pH 7.2, 12.5 mM MgCl_2_ and 200 μM UDP-GlcNAc (Sigma). To test *O*-fucosylation, the 50 μl reaction contained 10 μg *SPT-FLAG*, 5 μg *3TPR-SPY*, 50 mM Tris-HCl, pH 8.2, 50 mM NaCl, 5 mM MgCl_2_ and 200 μM GDP-fucose (Sigma). After incubation for 2 h at 25°C, the protein samples were separated by 10% SDS PAGE, and the band containing recombinant SPT (44 kDa) were excised and treated for MS analysis as described above.

### Mass spectrometry and data processing

Gel slices were prepared according to standard procedures adapted from Shevchenko et al. (2007)^59^. Briefly, the slices were washed with 50 mM TEAB buffer pH8 (Sigma), incubated with 10 mM DTT for 30 min at 65 °C followed by incubation with 30 mM iodoacetamide (IAA) at room temperature (both in 50 mM TEAB). After washing and dehydration with acetonitrile, the gels were soaked with 50 mM TEAB containing 10 ng/µl Sequencing Grade Trypsin (Promega) and incubated at 40 °C for 8 h. The extracted peptide solution was dried down, and the peptides dissolved in 0.1%TFA/3% acetonitrile. Aliquots were analysed by nanoLC-MS/MS on an Orbitrap Eclipse™ Tribrid™ mass spectrometer coupled to an UltiMate® 3000 RSLCnano LC system (Thermo Fisher Scientific, Hemel Hempstead, UK). The samples were loaded and trapped using a pre-column with 0.1% TFA at 15 µl min-1 for 4 min. The trap column was then switched in-line with the analytical column (nanoEase M/Z column, HSS C18 T3, 100 Å, 1.8 µm; Waters, Wilmslow, UK) for separation using the following gradient of solvents A (water, 0.1% formic acid) and B (80% acetonitrile, 0.1% formic acid) at a flow rate of 0.2 µl min-1: 0-3 min 3% B; 3-10 min increase B to 7% (curve 4); 10-70 min linear increase B to 37%; 70-90 min linear increase B to 55%; followed by a ramp to 99% B and re-equilibration to 3% B. Data were acquired with the following mass spectrometer settings in positive ion mode: MS1/OT: resolution 120K, profile mode, mass range m/z 300-1800, AGC 4e5, fill time 50 ms; MS2/IT: data dependent analysis with the following parameters: 1.5 s cycle time in IT turbo mode, centroid mode, isolation window 1 Da, charge states 2-5, threshold 1e4, HCD CE = 33, AGC target 1e4, max. inject time Auto, dynamic exclusion 1 count, 15 s exclusion, exclusion mass window ±10 ppm.

The raw data was processed in Proteome Discoverer 2.4 (Thermo Scientific, Waltham, USA). Spectra were recalibrated and identification was performed using an in-house Mascot Server 2.8.0 (Matrixscience, London, UK) with the TAIR10_pep_20101214 Arabidopsis thaliana protein sequence database (arabidopsis.org, 35,386 entries) to which the SPT-YFP fusion sequence was added. The Maxquant contaminants database (245 entries) was included in the search. Parameters were: enzyme trypsin, 2 missed cleavages, 6 ppm precursor tolerance, 0.6 Da fragment tolerance, carbamidomethylation (C) as fixed modification and oxidation (M), deamidation (N/Q), acetylation (protein N-terminus), dHex (NST, +146.058 Da), HexNAc (NST, +203.079 Da) as variable modifications. Evaluation was performed using Percolator. For peak detection and quantification, the Minora Feature Detector was used with a minimum trace length of 7 and S/N 3. Peptide abundances were determined as peak areas. After normalisation to total peptide amount the quantification was based on the top3 unique peptides per protein. Missing values were imputed by low abundance resampling. For hypothesis testing a background-based t-test was applied. Results were exported to Microsoft Excel. The percentage of the peptide modified with *O*-GlcNAc or *O*-fucose was calculated based on the normalised abundances of the modified peptide compared to the sum of the abundance of all versions of the peptide.

### Co-IP assay in tobacco leaves

*A. tumefaciens GV3101* strains harbouring the *35S::SPT-3xFLAG*, *35S::SEC-3HA* and *35S::SPY-3HA* constructs were transiently expressed either alone or co-infiltrated in 4-week old leaves of *N. benthamiana*. To enhance gene expression, an Agro strain harbouring P19 was always co-infiltrated. After 48 h from infiltration, the leaves were harvested and the total protein was extracted using the protein extraction buffer (25 mM Tris-HCl, pH 7.4, 1 mM EDTA, 150 mM NaCl, 10% glycerol, 0.15% NP-40, 1 mM NaF, 10 mM DTT, 2% PVPP, 1 mM PMSF and 1x protein inhibitor). Total protein extracts were immunoprecipitated with anti-HA magnetic beads (Thermo Scientific); input and immunoprecipitation (IP) samples were analysed by immunoblotting using anti-HA (Sigma) and anti-FLAG antibodies (Sigma) separately.

### Split-Luciferase Complementation (Split-Luc) Assay in tobacco leaves

The Split-Luc assay was performed as described previously^55^. *A.tumefaciens GV3101* strains harbouring the *35S::SEC-Cluc* and *35S::SPY-Cluc* constructs were co-infiltrated with either *35S::SPT-Nluc* or a *35S::Ø-Nluc* empty vector, while *35S::SPT-Nluc* was also co-infiltrated with *35S::Ø-Cluc* empty vector, into 4-week old leaves of *N.benthamiana*. To enhance gene expression, an Agro strain harbouring P19 was always co-infiltrated. After 48 h infection at room temperature, 0.4 mM D-luciferin [ThermoFisher] was infiltrated into the leaves, and LUC activity was measured using the NightOWL system equipped with a cooled CCD imaging apparatus (Berthold Technologies).

### Yeast-two-hybrid assay

For Yeast-two-Hybrid (Y2H) experiments coding DNA sequences were cloned into pDONR207 and recombined into *pGDAT7*® or *pGBKT7*® vectors (Clontech). Plasmids were transformed into the AH109 yeast strain by the lithium acetate (LiAc) method. Co-transformed strains were selected on SD/ -Leu-Trp (Sigma) at 28°C for 3-4 days. Transformed yeast were serially diluted (10^0^, 10^-1^, 10^-2^, 10^-3^) and dotted on SD/-Ade/-His/-Leu/-Trp and SD/-Ade/-His/-Leu/-Trp/ 2.5mM 3-amino-1,2,4-triazole (3-AT) to examine protein interactions. Growth was observed after -5 days of incubation at 28 °C.

### Chromatin Immunoprecipitation (ChIP)-qPCR assay

The ChIP assay was performed using young inflorescences (close sepals) of *SPT-sYFP spt-12* complementation line as previously described ^60^. Three biological repeats were performed, using 3 grams of plant material for each repeat. IP was conducted using the GFP-Trap beads (ChromTek). The enrichment of the *PINOID* promoter regions was quantified using quantitative PCR (qPCR) normalized with *ACTIN2* gene with the appropriate primers listed in Supplementary Table 1.

### Fluorescence resonance energy transfer (FRET)-Fluorescence-lifetime imaging microscopy (FLIM) assay

FRET-FLIM experiments were carried out using a Leica Stellaris 8 confocal microscope equipped with a 40x water immersion objective. Four-week-old *N. benthamiana* leaves were co-infiltrated with *GV3101* strains to express *SPT-EGFP* with either *RFP-NLS* or in combination with *IND-RFP*, *HEC1-RFP and SPT-RFP,* with or without *SEC-3xHA*, and *SPY-3xHA* recombinant proteins. To enhance gene expression, an Agro strain harbouring P19 was always co-infiltrated. After 48 h, fluorescence lifetime and efficiency were obtained with the supplied software (LAS X FLIM FCS). At least 60 nuclei per condition from three biological replicates were analysed.

### Transient transactivation assay

Transactivation assays were performed on 4-week old *N. benthamiana* Agro-infiltrated leaves using *pPID:GUS* alone, *pPID:GUS* combined with *p35S:SPT-GFP* and *p35S:IND-RFP* or together with either *p35S:SEC-HA, p35S:SEC-2-HA, p35S:SPY-HA* or *p35S:SPY-3-HA.* To enhance gene expression, an Agro strain harbouring P19 was always co-infiltrated. After 48 h from infiltration, leaves were harvested and the expression levels of *GUS* transcript were quantified via qRT-PCR, and the *hygromycin* gene in the *pPID:GUS* vector was used as internal control. Protein expression levels were detected by western blot. Primers used are listed in Supplementary Table 1.

### Western blot analysis

Protein samples were separated by 10% SDS-PAGE and transferred to a nitrocellulose membrane (GE Healthcare). After being blocked in 1 × PBST buffer containing 5% skimmed milk, the membrane was incubated with the selected primary antibody using a 1000-fold dilution overnight at 4 °C, washed three times with 1 × PBST (10 min each), and incubated with the selected secondary antibody conjugated with HRP using a 3000-fold dilution for 1 h at room temperature. After three washes with 1 × PBST (10 min each), the film was illuminated and photographed with ImageQuant800 (GE Healthcare). In case the primary antibodies were already conjugated with Horseradish Peroxidase (HRP), there was no need to incubate the membrane with secondary antibodies. The antibodies used were as following: anti-GFP (11814460001, Roche), anti-RFP (ab34771, Abcam), anti-FLAG (F3165, Sigma), anti-HA (3F10, Sigma), anti-mouse (sc-516102, Santa Cruz), anti-rabbit (ab205718, Abcam)

### Electrophoretic mobility shift assay (EMSA)

The EMSA reaction was performed as following: 150 ng amplified and gel-extracted wild-type or mutated DNA nucleotides together with indicated concentration of recombinant 6xHis-SPT protein were incubated in 1x EMSA buffer (250 mM Tris-HCl, pH 8.0, 500 mM NaCl, 25% glycerol, 10 mM DTT) on ice for 20 minutes. To test the effect of *O*-Glycosylation on SPT, recombinant SPT was firstly incubated with 5TPR-SEC or 3TPR-SPY recombinant proteins for 1 h at 25°C with 200 uM UDP-GlcNAc or GDP-fucose respectively, as previously described in the *In vitro* enzymatic assay section. Then the DNA nucleotides and EMSA buffer were added in the reaction. The reaction was analyzed by electrophoresis on 5% native acrylamide gel in 1x TBE buffer at 150 voltage for 50 minutes. After electrophoresis, the gels were stained with EB for 20 min followed with imaging by the UV imager (G:BOX F3 gel doc system, SYNGENE). Primers used for amplification of the wild-type and mutated *PID* 171-nt fragments were listed in Supplementary Table 1.

### Statistical analysis

Statistical analysis was performed as indicated in the figure legends, using Graphpad Prism9.

## Figure legends

**Extended Data Fig. 1.**
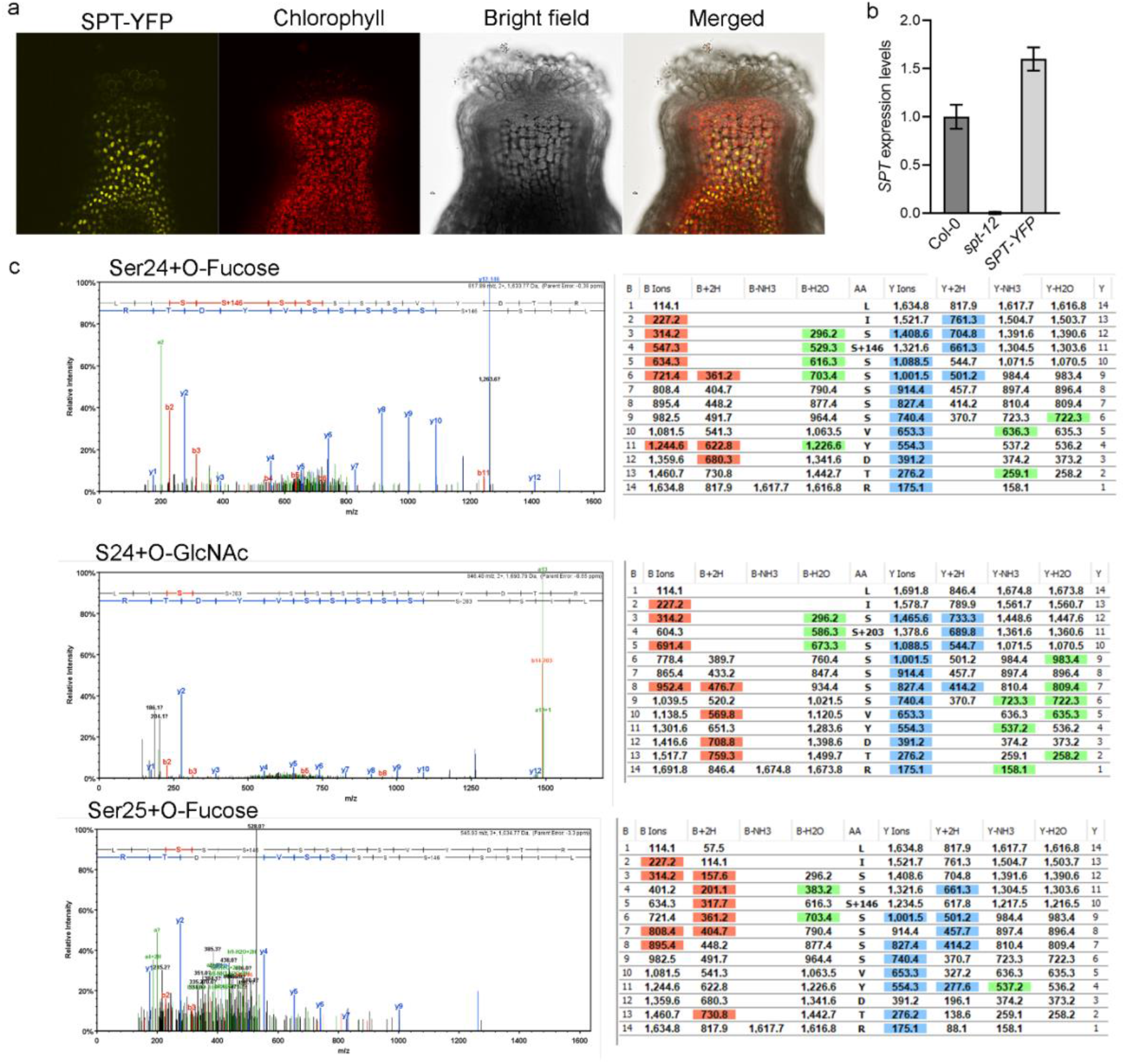
*In vivo* proteomic analysis of *SPT-YFP/spt* complementation line inflorescences. **a**, Confocal image (split-channels) of a stage-12 gynoecium apex of the *SPY-YFP/spt-12* complementation line, which fully restored the split-style phenotype of *spt-12* mutant. Note, expression of nuclear YPF signal is shown in the radial style. **b**, Quantification of *SPT* expression levels by qRT-PCR in inflorescences of Col, *spt-12* and *SPY-YFP/spt-12* complementation line. Values shown are means±SE. **c**, Representative spectra and ion tables showing modifications of SPT residue Ser24 by both *O-*fucose and *O-*GlcNAc (top and middle panels) as well as of residue Ser25 modified solely by *O-*fucose (lower panel).

**Extended Data Fig. 2.**
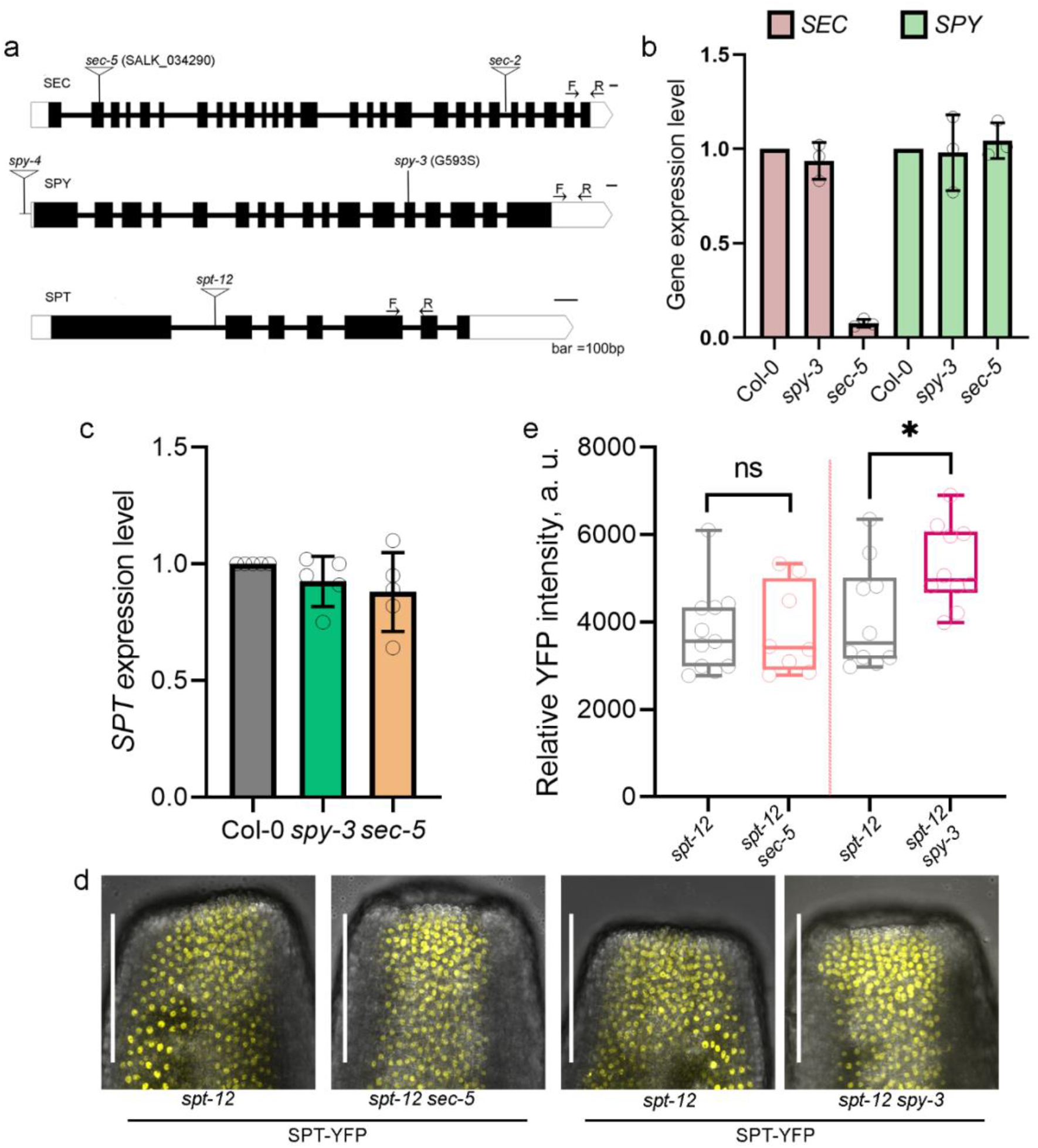
Quantification of *SPT*, *SPY*, and *SEC* expression patterns. **a**, Illustration of the SPT, SEC, and SPY gene structure indicating the T-DNA mutant lines and qRT-PCR primers used for these experiments. **b**, Quantification of *SEC* and *SPY* expression by qRT-PCR in mutant inflorescences of the transferase enzymes *sec-5* and *spy-3*. Values shown are means±SD from three biological repeats. **c**, Quantification of *SPT* expression by qRT-PCR in mutant inflorescences of the transferase enzymes *sec-5* and *spy-3*. Values shown are means±SD from three biological repeats. **d**, z-stack confocal images of stage-10 gynoecia of SPT complementation line in *sec-5* and *spy-3* backgrounds compared to their respective segregating control, *SPT-YFP/spt-12*. Scale bar represents 100 μm. **e**, Quantification of the YFP signal intensity in nuclei of gynoecia of genotypes depicted in panel d. Whiskers indicate min and max values. All data points are plotted in the graph. More than 10 individual gynoecia were used for each genotype. Significant differences are indicated in the graph following student’s *t* test. Ns, no significant difference. *, *P* < 0.05.

**Extended Data Fig. 3.**
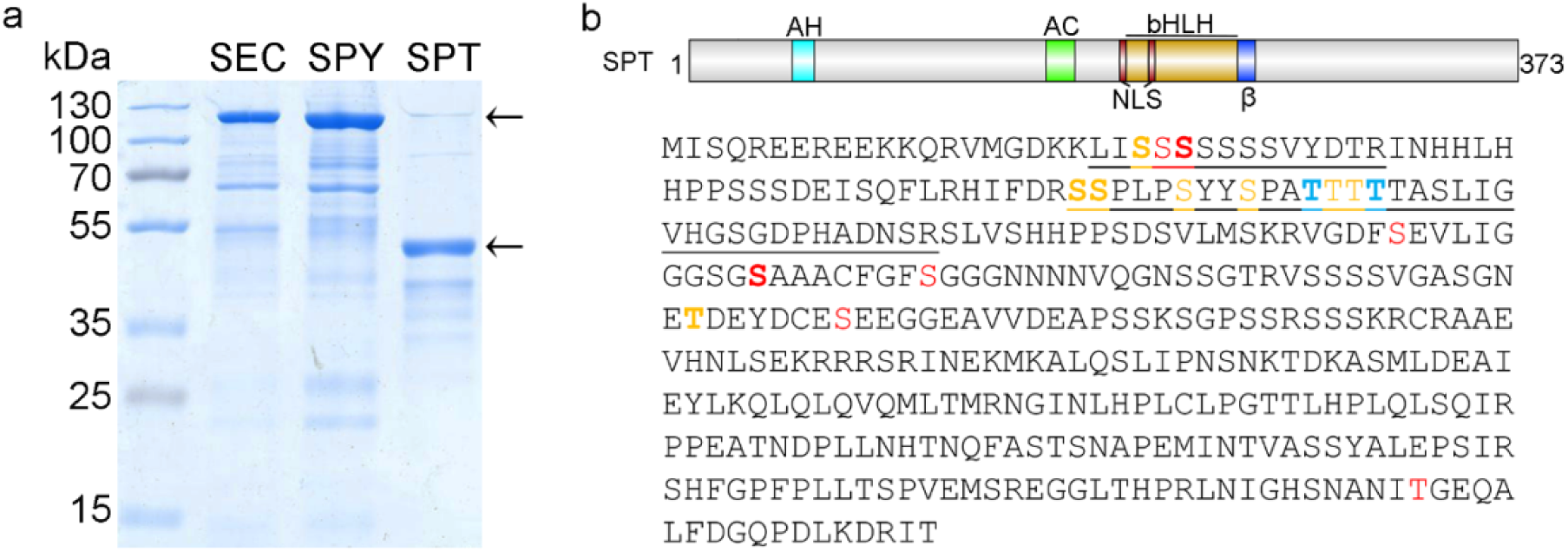
SPT is modified by SEC and SPY *in vitro*. **a**, SDS-PAGE gel showing purified SEC, SPY and SPT recombinant proteins from *E. coli*. The arrows indicate the band of the proteins at the expected size, *i.e*., *10xHis-MBP-5TPR-SEC* (SEC), 112 kDa; *10xHis-MBP-3TPR-SPY* (SPY), 107 kDa; and 6xHis-SPT (SPT), 44 kDa. **B**, Schematic representation of SPT proteins and summary of the residues found modified *in vitro* by *O-*GlcNAc (blue residues), *O-*fucose (red residues), and both modifications (yellow residues) along its amino acid sequence. S/T residues in bold were identified as frequently modified in the MS analysis. Peptide_1 and peptide_2 sequences are underlined.

**Extended Data Fig. 4.**
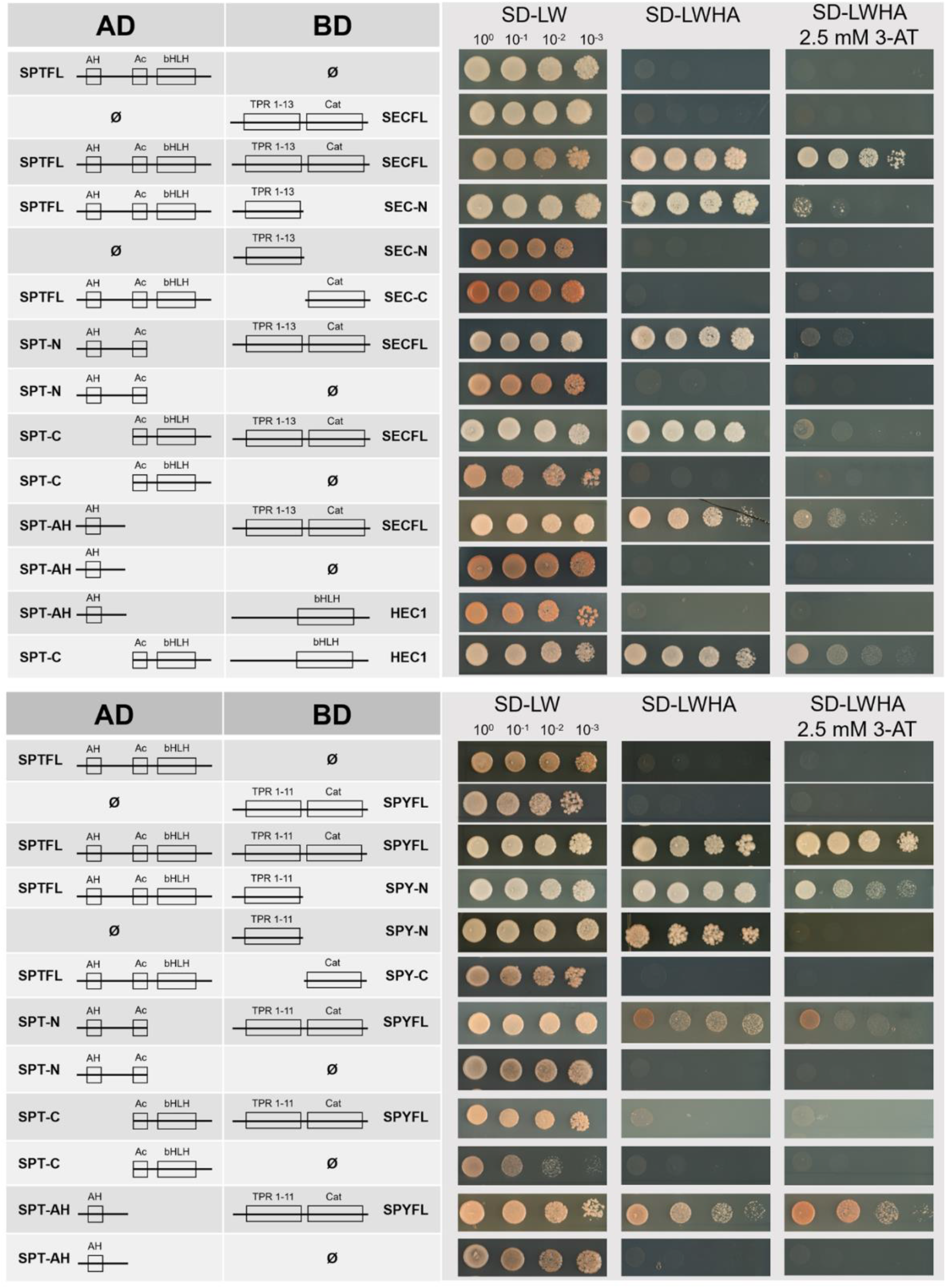
SEC and SPY interact with SPT via their N-terminal domains. Y2H experiments to determine SPT;SEC (top panel) and SPT;SPY (bottom panel) protein-protein interactions. Full-length proteins and functional domains (TPRs and Cat) structures are depicted on the left panel. HEC1 (top panel) was used as control to test interactions with SPT AH domain and the Ac plus bHLH domains combined. The assays were performed using SEC and SPY as the bait (BD) and SPT as prey (AD). Growth on selection media SD-LWHA supplemented with 2.5 mM 3-AT indicates strong interactors.

**Extended Data Fig. 5.**
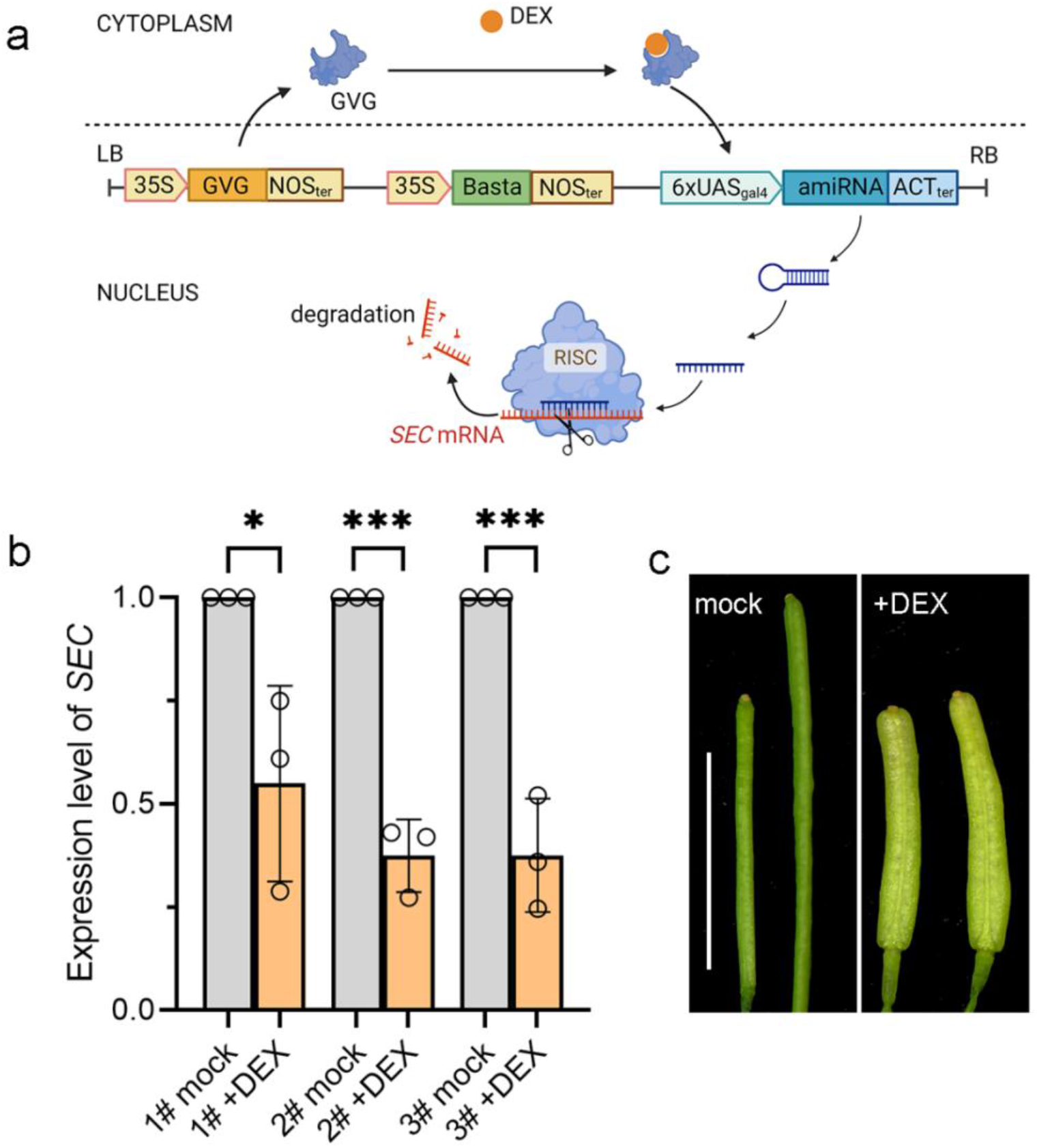
DEX-inducible RNAi of *SEC* inhibits fruit development in Arabidopsis. **a**, Schematic illustration of the working principle of the construct for DEX inducible RNAi of *SEC*. **b**, Quantification of *SEC* transcript levels by qRT-PCR from inflorescences of DEX-treated *spy-3 SEC RNAi* compared to mock. DEX treatment significantly downregulates *SEC* levels in three independent transgenic lines analysed. Values shown are means ± SD from three biological repeats. Significant differences are indicated in the graph following student’s *t* test. **c**, Early stages of fruit development after mock (left) and DEX (right) treatments of *spy-3 SEC RNAi* line. Scale bar indicates 1 cm.

**Extended Data Fig. 6.**
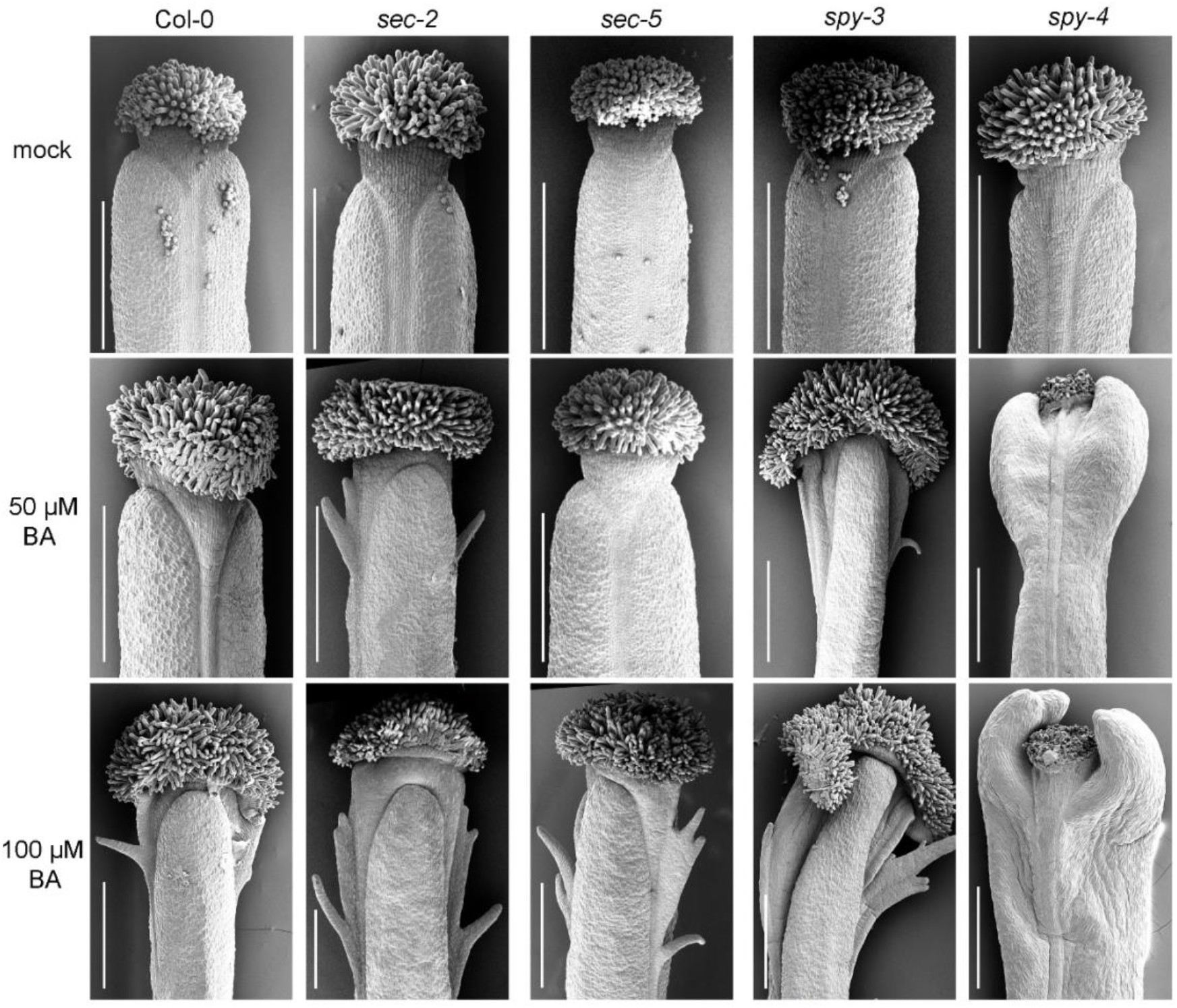
*spy* mutant gynoecia are hypersensitive to cytokinin treatments. Representative SEM images of stage-13 gynoecia of Col-0, *sec-2*, *sec-5*, *spy-3* and *spy-4* after pharmacological treatment with cytokinin (either 50 μM or 100 μM BA) and mock. Bars represent 500 μm.

**Extended Data Fig. 7.**
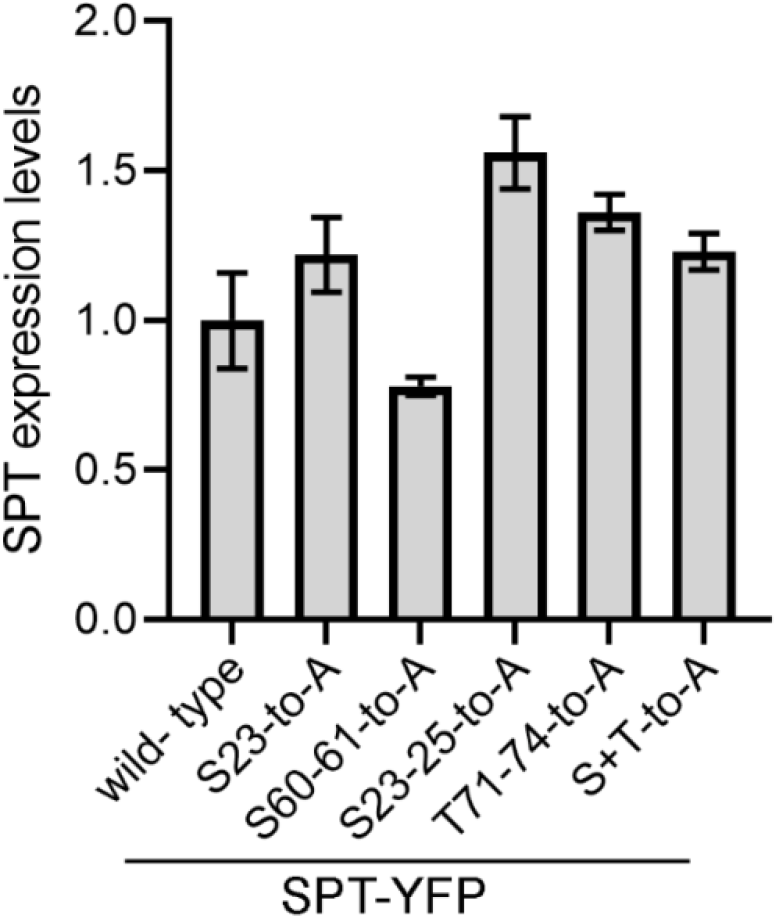
Molecular characterization of SPT-YFP complementation and point mutation lines. Quantification of *SPT* transcript levels in young inflorescences of SPT-YFP complementation and point mutation lines by qRT-PCR experiments. The experiment was performed once with four technical repeats. Values shown are means ± SE.

**Extended Data Fig. 8.**
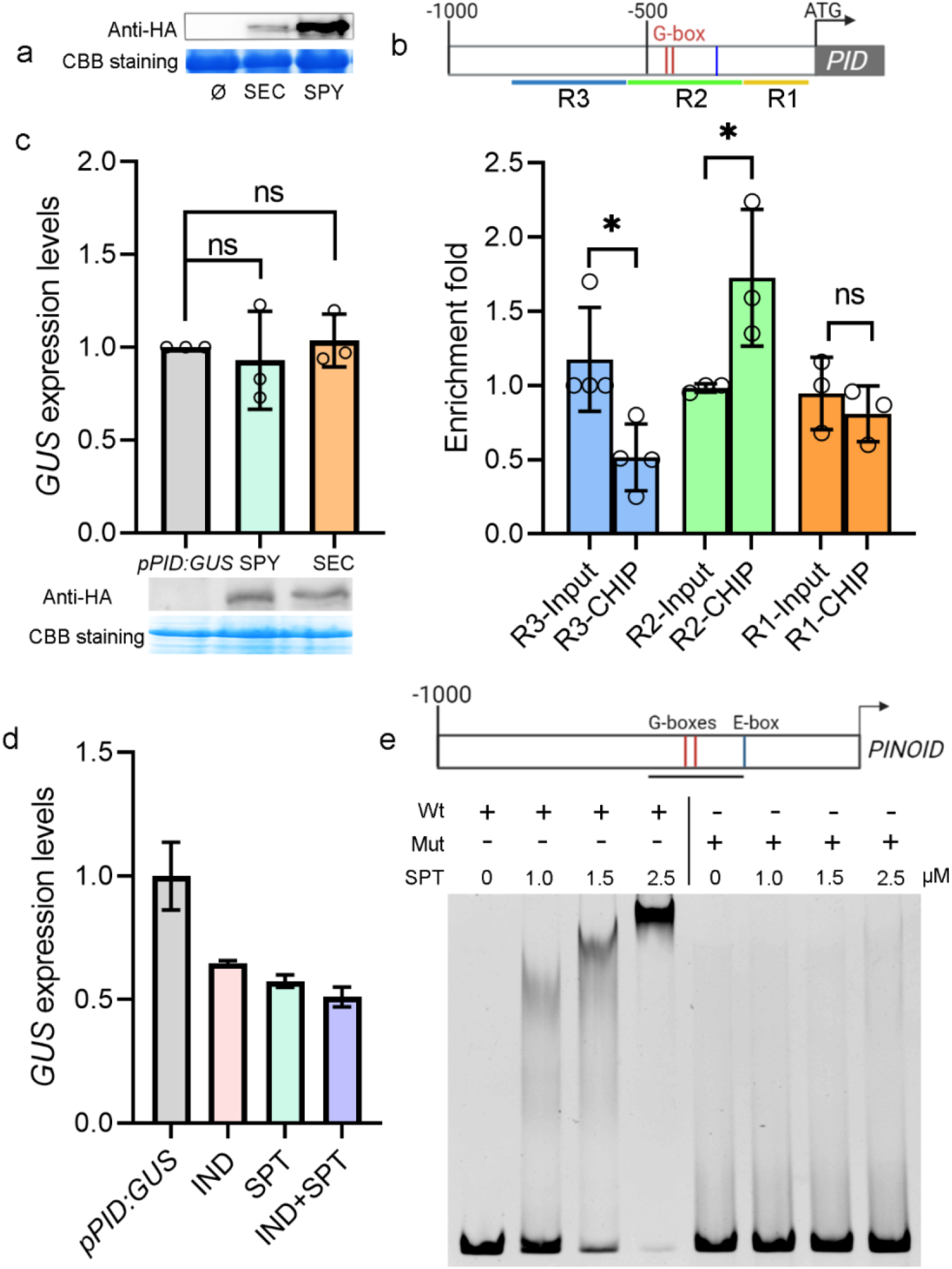
Molecular interaction between SPT and *PINOID* promoter *in vivo* and *in vitro*. **a**, Immunodetection of SEC-HA and SPY-HA in FRET-FLIM assay from *N. benthamiana* Agro-infiltrated leaves. The CBB (Coomassie Brilliant Blue) band was used as sample loading control. **b**, Top, schematic representation of the *PID* promoter including G-box (red line) and E-box (yellow line) and the position of the fragment amplified across those *cis-* elements. Bottom, qRT-PCR quantification of ChIP experiments carried out from young inflorescences of *SPT-YFP/spt-12* complementation line. The enrichment of the *SPT* binding to the region (R2) containing the two G-boxes demonstrate SPT binds *in vivo* to the *PID* promoter. Values shown are means±SD from three biological repeats. Significant differences are indicated in the graph following student’s *t* test. **c**, Top, qRT-PCR quantification of *GUS* expression in transactivation assay from tobacco leaves infiltrated with either the *pPID:GUS* construct or SEC:HA (*p35S:SEC-HA*) or SPY:HA (*p35S:SPY-HA*) .The OD values here used for both SEC and SPY were 0.5 in each experiment. Note, SEC and SPY have no effect on p*PID:GUS* expression. Values shown are means ± SD from three repeats. ns, no significant difference by student’s *t* test. Bottom, immunodetection of SEC-HA and SPY-HA from *N. benthamiana* Agro-infiltrated leaves. The CBB (Coomassie Brilliant Blue) band was used as sample loading control. **d**, qRT-PCR experiments showing quantification of *GUS* expression in transactivation assay from tobacco leaves following infiltration with IND and SPT alone or both compared to the *pPID:GUS* construct alone. The OD values used for both SPT and IND were 0.5 in each experiment. Values shown are means ± SE. **e**, graphic representation of the PID promoter including the 171-bp fragment used in EMSA experiments. Recombinant SPT (6xHis-SPT) binds to wild-type sequences of the G-box (red lines) present in the *PINOID* promoter, in a concentration-dependent manner (left panel). Incubation with a mutated (Mut) G-boxes (from CACGTG to TGATGA) abolish this binding. Similar results were obtained from two independent experiments.

## Supplementary information

**Supplementary table 1.** Primers used in this study.

**Supplementary dataset1.** Mass Spectrometry proteomics data have been deposited to the ProteomeXchange Consortium via the PRIDE partner repository with the dataset identifier PXD037917 (https://www.ebi.ac.uk/pride/archive/projects/PXD037917/private).

**Supplementary dataset2. All original images of western blotting used in this study.**

